# A Small-Molecule Activity-Based Probe for Monitoring Ubiquitin C-terminal Hydrolase L1 (UCHL1) Activity in Live Cells and Zebrafish Embryos

**DOI:** 10.1101/827642

**Authors:** Paul P. Geurink, Raymond Kooij, Aysegul Sapmaz, Sijia Liu, Bo-Tao Xin, George M. C. Janssen, Peter A. van Veelen, Peter ten Dijke, Huib Ovaa

## Abstract

Many reagents have been emerged to study the function of specific enzymes *in vitro*. On the other hand, target specific reagents are scarce or need improvement allowing investigations of the function of individual enzymes in a cellular context. We here report the development of a target-selective fluorescent small-molecule activity-based DUB probe that is active in live cells and whole animals. The probe labels active Ubiquitin Carboxy-terminal Hydrolase L1 (UCHL1), also known as neuron-specific protein PGP9.5 (PGP9.5) and parkinson disease 5 (PARK5), a DUB active in neurons that constitutes 1-2% of total brain protein. UCHL1 variants have been linked with the neurodegenerative disorders Parkinson’s and Alzheimer’s disease. In addition, high levels of UCHL1 also correlate often with cancer and especially metastasis. The function of UCHL1 or its role in cancer and neurodegenerative disease is poorly understood and few UCHL1 specific research tools exist. We show that the reagents reported here are specific for UCHL1 over all other DUBs detectable by competitive activity-based protein profiling and by mass spectrometry. Our probe, which contains a cyanimide reactive moiety, binds to the active-site cysteine residue of UCHL1 irreversibly in an activity-dependent manner. Its use is demonstrated by labelling of UCHL1 both *in vitro* and in cells. We furthermore show that this probe can report UCHL1 activity during the development of zebrafish embryos.

## INTRODUCTION

The Ubiquitin system relies to a great extent on cysteine catalysis. Ubiquitin is a small protein that consists of 76 amino acids that can modify target proteins through lysine residues although it is also occasionally found to modify N-termini as well as cysteine and threonine residues.^1-3^ Addition of ubiquitin is catalyzed by E1 (2), E2 (∼40) and E3 (>600) enzymes in an ATP-dependent conjugation reaction by specific combinations of E1, E2 and E3 enzymes and it is reversed by any of ∼100 deubiquitylating enzymes (DUBs) in humans.^4, 5^ The enzyme Ubiquitin Carboxy-terminal Hydrolase L1 (UCHL1), also known as neuron-specific protein PGP9.5 (PGP9.5) and parkinson disease 5 (PARK5), is a small protease that is thought to remove ubiquitin from small substrates and it belongs to the small family of Ubiquitin C-terminal Hydrolases (UCHs).^6^

It is clear that UCHL1 can cleave ubiquitin and that mutation and reduced activity of this enzyme have been associated with neurodegenerative diseases, including Parkinson’s and Alzheimer’s disease.^7-12^ High UCHL1 levels correlate with malignancy and metastasis in many cancers^13, 14^ and have also been attributed to cellular stress, although the molecular mechanism of all these processes is unclear.

We earlier observed extreme levels of UCHL1 activity in lysates from prostate and lung cancer cells using a ubiquitin-derived activity-based probe that targets all cysteine DUBs.^15^ We reasoned that a good cell-permeable activity-based probe that targets UCHL1 specifically amongst other cysteine DUBs would be a highly valuable tool to understand its function in malignant transformation and its role in the development of neurodegenerative diseases.

UCHL1, like many DUBs, is a cysteine protease, a class of enzymes considered extremely difficult to inhibit with small molecules as this class of enzymes is associated with unspecific reaction with cysteine alkylating agents and with redox-cycling artifacts in assays.^16^ In addition, DUBs intrinsically bind ubiquitin through a protein-protein interaction, which is by definition difficult to interfere with using small molecules. Many DUBs, including UCHL1, are inactive without a substrate and substrate binding aligns the catalytic triad for cleavage.^17^ Nevertheless, recently significant successes have been booked in the development of reversible and irreversible selective small-molecule inhibitors of the DUB USP7.^18-23^ We have recently reported the development of a selective covalent small-molecule inhibitor of the DUB ovarian tumor (OTU) protease OTUB2 using a covalent fragment approach and parallel X-ray crystallography.^24^ We reasoned that such covalent molecules are a good inroad for the further elaboration of specific activity-based probes (ABPs) also inspired by earlier work from the Tate lab that recently reported a small-molecule broadly acting DUB probe.^25^ We were pleased to find a good starting point in patent literature^26^ that we used in our studies for the design of fluorescent ABPs. We here report the development of a fluorescent small-molecule ABP that can report UCHL1 activity in human cells and in zebrafish embryos.

## RESULTS AND DISCUSSION

The development of a small-molecule-based DUB ABP starts with the identification of an appropriate DUB-selective small-molecule covalent binder. We reasoned that an ideal compound needed to meet two criteria: 1) it binds covalently to the active-site cysteine residue of a DUB and 2) it can easily be modified by chemical synthesis. Our attention was drawn to a collection of (*S*)-1-cyanopyrrolidine-3-carboxamide-based compounds reported to inhibit UCHL1 activity with submicromolar affinity.^26^ These compounds are equipped with a cyanimide moiety that is known to react with thiols to form an isothiourea covalent adduct (Figure 1A) and thought to react reversibly.^27^ Despite the expected reversible nature we decided to investigate this compound as a potential probe starting point.

**Figure 1.**
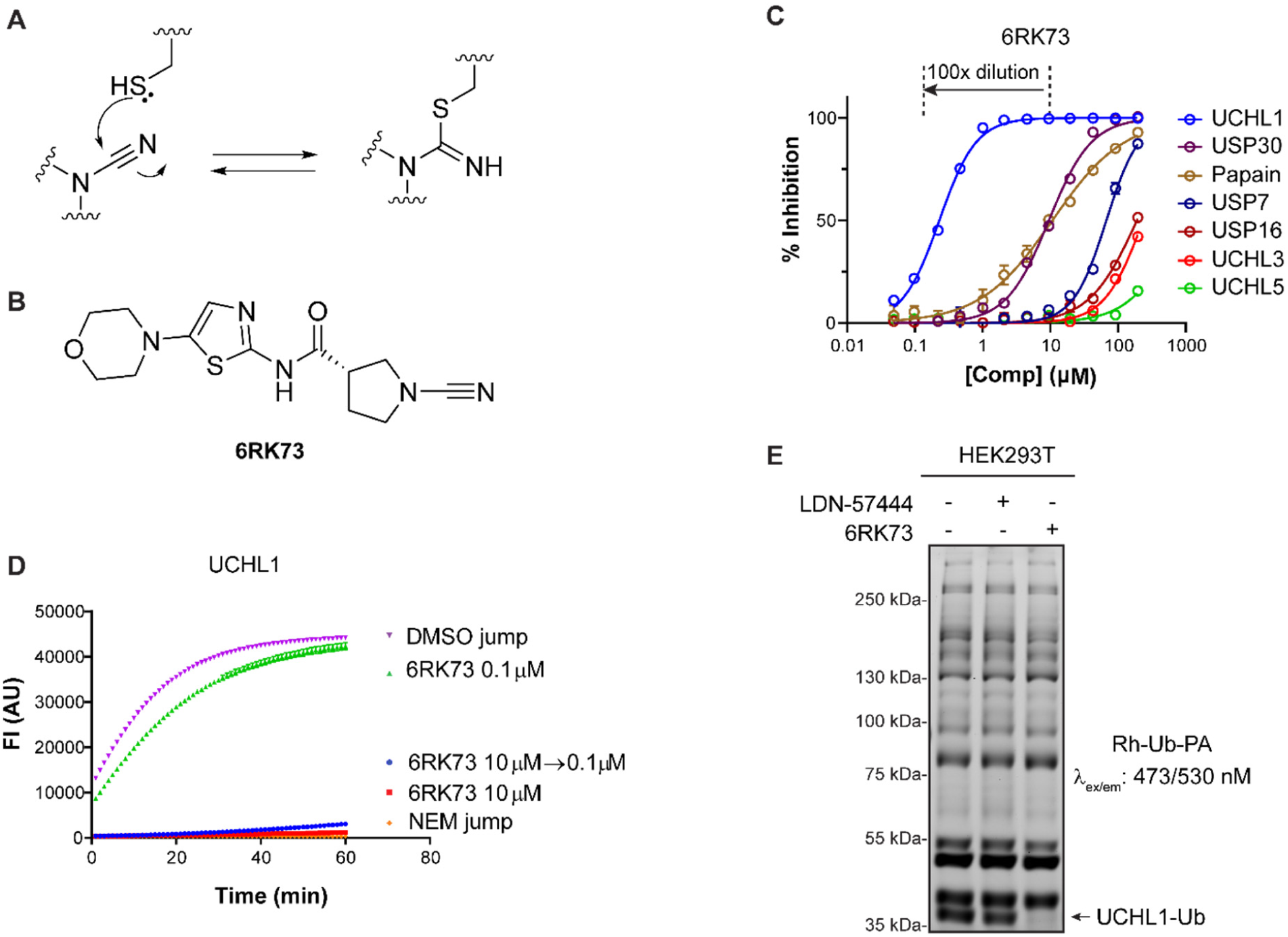
Biochemical characterization of UCHL1 inhibitor **6RK73**. A) Reaction of a thiol with a cyanimide results in the formation of an isothiourea adduct. B) Structure of UCHL1 inhibitor **6RK73**. C) IC_50_ determination of **6RK73** for UCHL1, UCHL3 and UCHL5. D) Progress curves for UCHL1 proteolytic activity after jump dilution (see also Figure C). DMSO and *N*-ethylmaleimide (NEM) are used as controls. E) Deconvoluted mass spectra of UCHL1 before (blue) and after (red) reaction with **6RK73**. F) Fluorescence labelling of remaining DUB activity in HEK293T cells upon treatment with UCHL1 inhibitors LDN-57444 and **6RK73**.

### Characterizing UCHL1 cyanimide inhibitors

In order to gain insight into the mode of action and DUB selectivity of these inhibitors we synthesized and characterized one compound (compound **6RK73**, Figure 1B) that in our hands inhibits UCHL1 with an IC_50_ of 0.23 µM after 30 minutes of incubation in a biochemical activity assay using fluorogenic Ub-Rho-morpholine^28^ substrate (for preparation see Supporting Information) in the presence of 2 mM cysteine. Beneficially, **6RK73** proved to be almost unreactive towards the closest DUB family members UCHL3 and UCHL5 (Figure 1C). Selectivity for UCHL1 was further confirmed by IC_50_ determination against a panel of other cysteine DUBs (including USP7, USP30 and USP16) and the non-DUB cysteine protease papain, showing over 50-fold difference in IC_50_ value (Figure 1C and Supporting Information Table S1). We next performed a jump dilution experiment^29^ in which 100× final assay concentration of UCHL1 was treated with 10 µM of **6RK73** followed by 100 times dilution into substrate-containing buffer and direct fluorescence read-out (Figure 1C, D). Only after 30 minutes a negligible increase in fluorescence signal could be detected which indicates that the inhibitor acts basically irreversible. The formation of a stable covalent complex between UCHL1 and a single **6RK73** molecule was confirmed in an experiment where UCHL1 was incubated with **6RK73** and the reaction followed by LC-MS analysis (see Supporting Information). Next, we investigated whether the compound would inhibit UCHL1 in live cells. HEK293T cells were treated with 5 µM **6RK73** or the commercially available active-site directed reversible UCHL1 inhibitor LDN-57444^30^ for 24h, followed by cell lysis and treatment with the fluorescent broad-spectrum DUB probe Rhodamine-Ubiquitin-propargylamide (Rh-Ub-PA) to label all residual cysteine-DUB activity.^31, 32^ The samples were denatured, resolved by SDS-PAGE and scanned for Rhodamine fluorescence (Figure 1E). Each band represents an active DUB that reacted with the probe and the ability of a compound to inhibit a DUB is reflected by disappearance of its corresponding band. Indeed, the band belonging to UCHL1^33^ disappears upon treatment with **6RK73**, whereas all other bands remain unchanged, indicating that **6RK73** selectively inhibits UCHL1 in the presence of other DUBs in cells. In comparison, UCHL1 is hardly inhibited by LDN-57444 in this experiment, despite their comparable IC_50_ values (0.88 µM for LDN-57444), which might be attributed to the fast-reversible nature of this inhibitor.^30^

### From inhibitor to probe

Given the high inhibitory potency and UCHL1 selectivity both *in vitro* and in cells and the fact that it forms an irreversible covalent bond we envisioned that this type of cyanimide-containing molecules can serve as an ideal starting point for the construction of small-molecule selective DUB ABPs. This would require the instalment of a reporter group (e.g. fluorescent label) onto the molecule. Upon close inspection of **6RK73** however, we realized that this molecule does not provide an appropriate site for modification. We therefore generated azide **8RK64** to which then several reporter groups were coupled using the copper(I)-catalyzed azide alkyne cycloaddition (CuAAC) or ‘click reaction’. The compounds and their synthesis routes are shown in Scheme 1. Compound **2** was synthesized from 4-piperidinone (**1**) in four steps according to a reported procedure.^26^ The Fmoc-protected piperidine amine was liberated with DBU and coupled to 2-azidoacetic acid resulting in compound **3**. Next, the Boc protecting group was removed from the pyrrolidine amine, followed by a reaction with cyanogen bromide to install the cyanimide moiety resulting in **8RK64**. Treatment of UCHL1 with this compound followed by IC_50_ determination and LC-MS analysis gave results comparable to those for **6RK73** (Figure 2A, B, Supporting Information), which indicates that **8RK64** also functions as a UCHL1 covalent inhibitor. With an IC_50_ value of 0.32 µM towards UCHL1 and 216 µM and >1 mM towards UCHL3 and UCHL5 respectively (Figure 2A, Supporting Information Table S1), this compound also retained its UCHL1 selectivity. In addition, **8RK64**, like **6RK73**, also inhibits UCHL1 activity in cells as shown in a DUB profiling experiment in HEK293T cells using a Cy5-Ub-PA probe (Figure 2C). Notably, **8RK64** could potentially be used as ‘2-step ABP’ by taking advantage of its azide moiety.^34^

**Figure 2.**
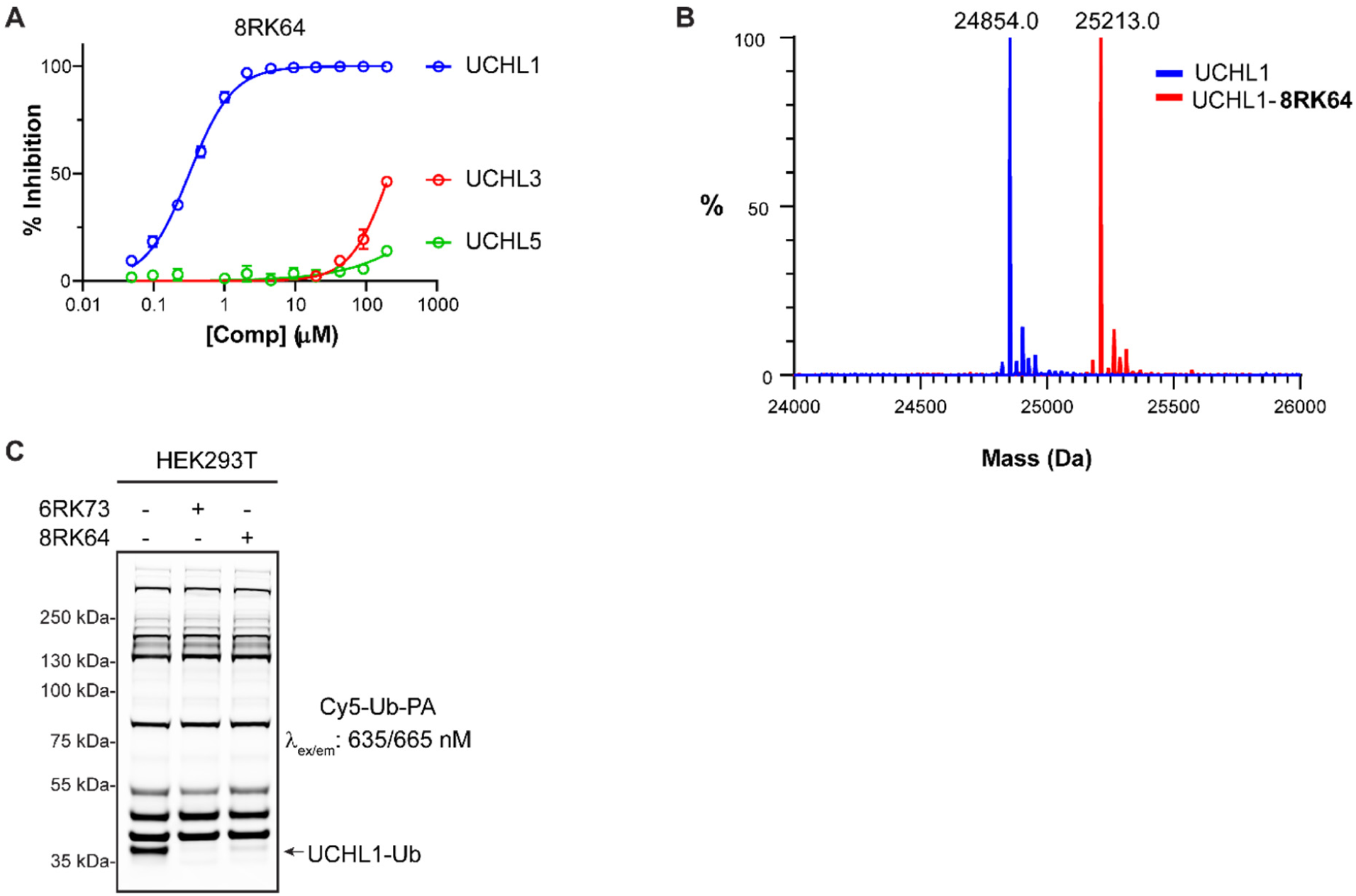
Biochemical characterization of **8RK64**. A) IC_50_ determination of **8RK64** for UCHL1, UCHL3 and UCHL5. B) Deconvoluted mass spectra of UCHL1 before (blue) and after (red) reaction with **8RK64**. C) Fluorescence labelling of remaining DUB activity in HEK293T cells upon treatment with UCHL1 inhibitors **8RK64** and **6RK73**.

### Installation of a dye preserves inhibitory properties

As it was unclear what the effect of coupling a bulky fluorescent group would have on the UCHL1 inhibition profiles and cell permeability we decided to test three commonly used fluorophores. BodipyFL-alkyne, BodipyTMR-alkyne^35^ and Rhodamine110-alkyne (for preparation see Supporting Information) were coupled using copper(I)-mediated click chemistry to the azide of **8RK64**, resulting in compounds **8RK59, 9RK15** and **9RK87** (Scheme 1). These ‘one-step’ ABPs can potentially be used for visualization of UCHL1 activity without the need for additional bio-orthogonal chemistry procedures. IC_50_ determination of these probes against UCHL1 revealed that the instalment of the dyes affected the inhibitory potency only marginally (Figure 3A and Supporting Information Table S1). Rhodamine110 probe **9RK87** is almost as potent as its azide precursor **8RK64** with IC_50_ values of 0.44 µM and 0.32 µM respectively. Instalment of BodipyTMR (**9RK15**) on the other hand, resulted in a 10-fold potency decrease, although the data points could not be fitted properly to a dose-response function. The less bulky BodipyFL-ABP **8RK59**, although not as potent as **8RK64**, showed a very acceptable inhibition of UCHL1 with an IC_50_ close to 1 µM. The ability of **8RK59** to form a covalent complex with UCHL1 was confirmed in an LC-MS experiment as described above (Supporting Information).

**Figure 3.**
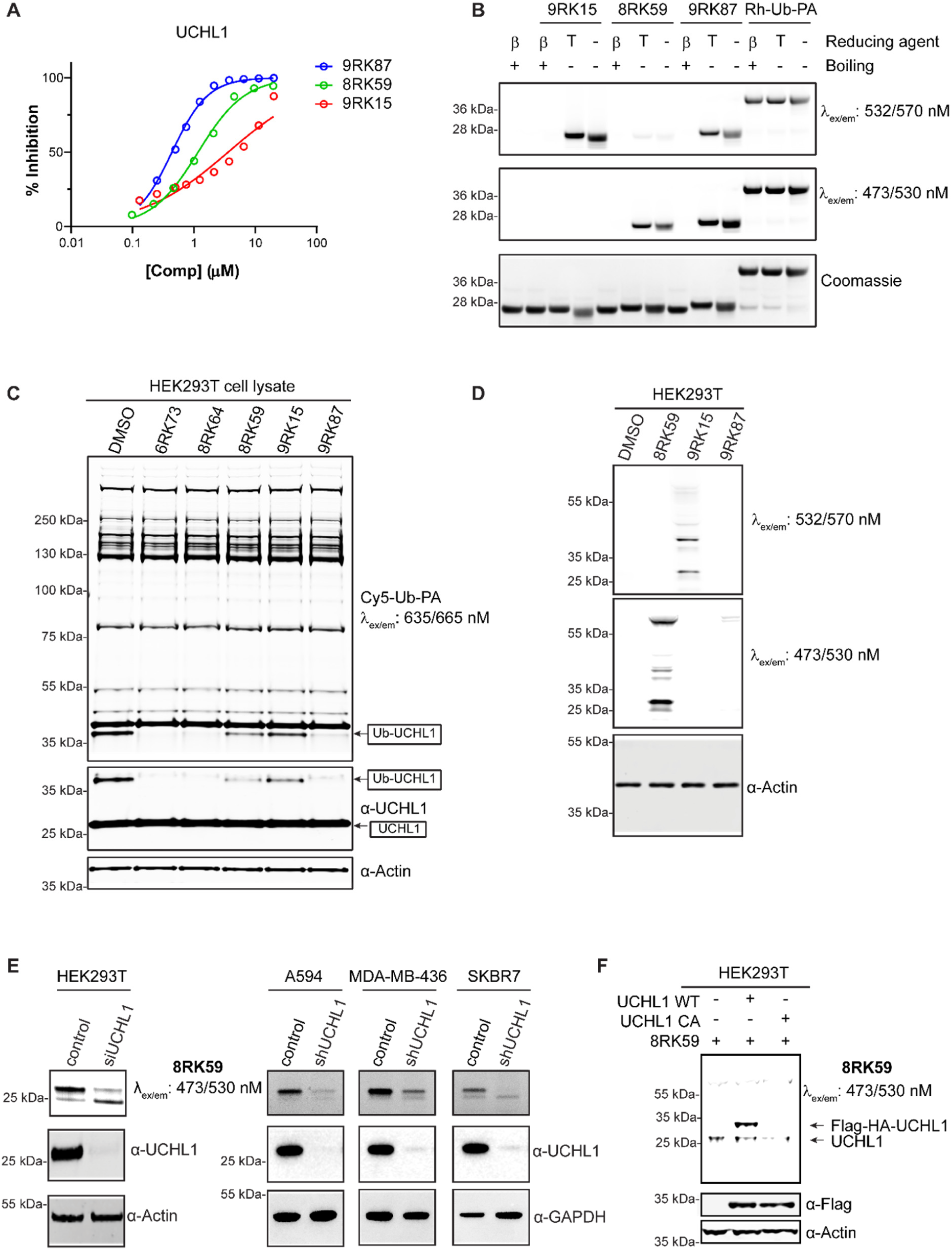
Characterization of the fluorescent UCHL1 probes *in vitro* and in cells. A) IC_50_ determination of **8RK59, 9RK15** and **9RK87** for UCHL1. B) Labelling of purified recombinant human UCHL1 by the three probes and Rh-Ub-PA. β: β-mercaptoethanol; T: TCEP. C) Fluorescence labelling by Cy5-Ub-PA of remaining DUB activity in HEK293T cell lysate upon treatment with UCHL1 inhibitors and probes. D) Fluorescence scans showing the labelling pattern in HEK293T cells of the three probes. E) Fluorescence labelling of UCHL1 activity in HEK293T, A549, MDA-MB-436 and SKBR7 cells with **8RK59**. F) **8RK59** labels overexpressed Flag-HA-UCHL1 wt but not the C90A active site mutant in HEK293T cells.

### ABPs can visualize UCHL1 activity and the covalent linkage is thermally reversed

We next set out to investigate whether the probes can be used to label and visualize UCHL1 activity after SDS-PAGE and fluorescence gel scanning similar to the Rh-Ub-PA probe. To our surprise for none of the three small-molecule probes a clear band corresponding to probe-labelled UCHL1 could be detected after incubation with purified recombinant human UCHL1. We reasoned that the isothiourea bond between UCHL1 and probe, which is stable under the conditions used for inhibition and LC-MS experiments (*vide supra*), might be susceptible to the conditions used for protein denaturation, e.g. boiling in the presence of ∼300 mM β-mercaptoethanol. Indeed, when the same samples were resolved by SDS-PAGE under non-denaturing conditions (no boiling and absence of β-mercaptoethanol) a clear band appeared that corresponds to probe-labelled UCHL1 for all three probes (Figure 3B). We also investigated if the ABP-UCHL1 bond would survive when β-mercaptoethanol is replaced by tris(2-carboxyethyl)phosphine) (TCEP), both of which are used to create a reducing environment. Figure 3B clearly shows that the ABP-UCHL1 bands remain intact in the presence of 50 mM TCEP and show a better-resolved profile (less smearing) compared to the non-reducing samples. The Rh-Ub-PA control samples show that nearly all UCHL1 is labeled and that the formed bond for this probe is stable under denaturing conditions, which corroborates earlier findings.^31^ The bands corresponding to Rh-Ub-PA and **9RK87** bound to UCHL1 (both bearing the same dye and present in equal amounts) are of similar intensity, which indicates that the small-molecule probes bind UCHL1 efficiently and that all UCHL1 is active upon probe engagement.

### ABPs bind to the active site cysteine residue of UCHL1 and visualize UCHL1 activity in various cell lines

We next assessed the ability of the probes to bind and inhibit UCHL1 in a cell lysate by treating HEK293T cell extracts with 5 µM of the three fluorescent probes, as well as their azide precursor **8RK64** and inhibitor **6RK73** for 1 hour followed by labelling of all residual DUB activity with Cy5-Ub-PA. The Cy5-labelled Ub probe was used here to circumvent spectral interference with either of the other dyes used in the small-molecule probes. Fluorescent scanning of the gel after SDS-PAGE as well as Western blotting using anti-UCHL1 antibody clearly showed that Rhodamine probe **9RK87** inhibits UCHL1 activity similar to **8RK64** and **6RK73** (Figure 3C). Both Bodipy probes also potently inhibit UCHL1 in a cell lysate, although to a somewhat lesser extent, which could be expected on the basis of their IC_50_ values. All other bands are unchanged, which demonstrates that all compounds are able to bind UCHL1 selectively with respect to other DUBs in a cell lysate.

Encouraged by these results we set out to assess the ability of the probes to penetrate the cell membrane and to label active UCHL1 in cells. HEK293T cells were treated with 5 µM of the probes for 24 hours followed by cell lysis, SDS-PAGE (in the absence of β-mercaptoethanol and boiling) and fluorescence scanning at two wavelengths to detect all fluorescent dyes (Figure 3D). A clear band just above 25 kDa is observed for both Bodipy probes (**8RK59** and **9RK15**), which likely corresponds to ABP-labelled UCHL1 with an expected mass of ∼25.5 kDa. In addition to this band a few extra bands are visible including one just below UCHL1 and one more pronounced band around 55 kDa. Interestingly, hardly any band can be seen for the so-far most potent probe **9RK87**. We attributed this effect to the difference in cell permeability between Bodipy and Rhodamine dyes, with the latter known to be less capable of crossing the cell membrane.^36^ Indeed, upon further investigation using microscopy in ABP-treated HeLa and HEK293T cells we confirmed that Rhodamine probe **9RK87** is unable to enter these cells, whereas both Bodipy ABPs clearly are (Supporting Information Figure S1). For this reason and because the BodipyFL-ABP proved to be a better inhibitor compared to its BodipyTMR analogue we decided to continue with **8RK59** as the preferred probe for all further experiments.

The ability of **8RK59** to label UCHL1 activity in different cell lines was further explored in HEK293T cells and in three cancer cell lines known to express high levels of endogenous UCHL1: non-small cell lung cancer (NSCLC) A549 cells, triple negative breast cancer (TNBC) MDA-MB-436 cells and SKBR7 cells.^37^ Cells transfected with UCHL1 shRNA knock-down (shUCHL1) or siUCHL1 as well as empty vector control or scrambled oligo (siControl) were treated with 5 µM of each probe for 24 hours, followed by cell lysis, SDS-PAGE (without boiling and β-mercaptoethanol) and fluorescence scanning (Figure 3E). A clear band appears in the fluorescence scan at the expected height (∼25.5 kDa) in all four cell lines and this band is significantly decreased in the UCHL1 knock-down samples, indicating that this band indeed corresponds to ABP-labelled UCHL1.

To confirm that **8RK59** binds the active site cysteine residue in UCHL1 we overexpressed Flag-HA-tagged UCHL1 and its C90A catalytic inactive mutant in HEK293T cells and incubated these cells with 5 µM **8RK59** for 24 hours. Fluorescence scanning and anti-FLAG Western blotting shows that **8RK59** only binds to wild-type UCHL1 but not to catalytically inactive UCHL1, indicating that the probe binding site is the active site cysteine (Figure 3F).

### Determination of DUB selectivity and potential off-targets of the ABP

As mentioned before, besides the band corresponding to ABP-labelled UCHL1 a few other bands appeared on gel (Figure 3D) but based on the DUB profiling results (Figure 3C) these bands can most likely not be attributed to other DUBs. In order to gain more insight into potential off-targets we performed pull-down experiments coupled to mass spectrometry to identify the proteins binding to our probe. We started with a ‘2-step ABP’ approach in which HEK293T cells were incubated with azide-containing compound **8RK64** or DMSO control, followed by a post-lysis click reaction with biotin-alkyne^38^ and subsequent pull-down with neutravidin-coated beads (Supporting Information Figure S2A, B). Samples were run (1 cm) on a SDS-PAGE gel, lanes were cut into two pieces and the proteins were subjected to trypsin digestion and analyzed by LC-MSMS. As expected, the most enriched protein identified from this experiment was UCHL1 (Supporting Information Figure S2C). Only one additional protein was also highly enriched, a protein deglycase named DJ-1 (PARK7) with a molecular weight of 20 kDa, which most likely corresponds to the band just below UCHL1 in Figure 3D. This enzyme also harbors an active site cysteine residue which could potentially bind to our probe. Indeed, incubation of UCHL1 and PARK7 knock-down cells with **8RK59**, followed by anti-UCHL1 and anti-PARK7 Western blotting, revealed that PARK7 also reacts with **8RK59** and that the gel band just below UCHL1 corresponds to PARK7 (Supporting Information Figure S2D).

In addition to UCHL1 and PARK7, a few other bands can be seen on gel, yet we only identified these two enzymes in the 2-step ABP approach. We therefore performed a 1-step pull-down experiment where we used two biotinylated versions of **8RK64**: compound **11RK72** where biotin is directly linked to the inhibitor and compound **11RK73** with a PEG spacer in between. Both compounds show high inhibitory potential towards UCHL1 (Figure 4A) and form a covalent bond with UCHL1 (Supporting Information). HEK293T cell lysate was incubated with both biotin-ABPs, followed by pull-down with neutravidin-coated beads and subjected to full proteome LC-MSMS analysis (Figure 4B, Supporting Information Figure S2E). Efficient UCHL1 pull-down was confirmed for both biotinylated probes but not the DMSO and biotin-alkyne-treated control samples by Western blotting using anti-UCHL1 antibody (Figure 4C). From the LC-MSMS data, the relative protein abundances were calculated in the pull-down samples and compared to control samples. The list of identified proteins was ranked for total abundance to identify the highest enriched proteins (Supporting Information). Inspection of the list of all enzymes related to Ub (Ub-like proteins, DUBs, E1, E2 and E3 ligases) further substantiates the specificity of the probes for UCHL1 within the Ub system as shown in Figure 4D. Only a few of these enzymes were identified in the pull-down experiment with at least 150-fold lower abundance compared to UCHL1. The first other DUB on the list is UCHL3, the closest UCHL1 family member, and one of the most abundant DUBs in cells, which could explain this result.

**Figure 4.**
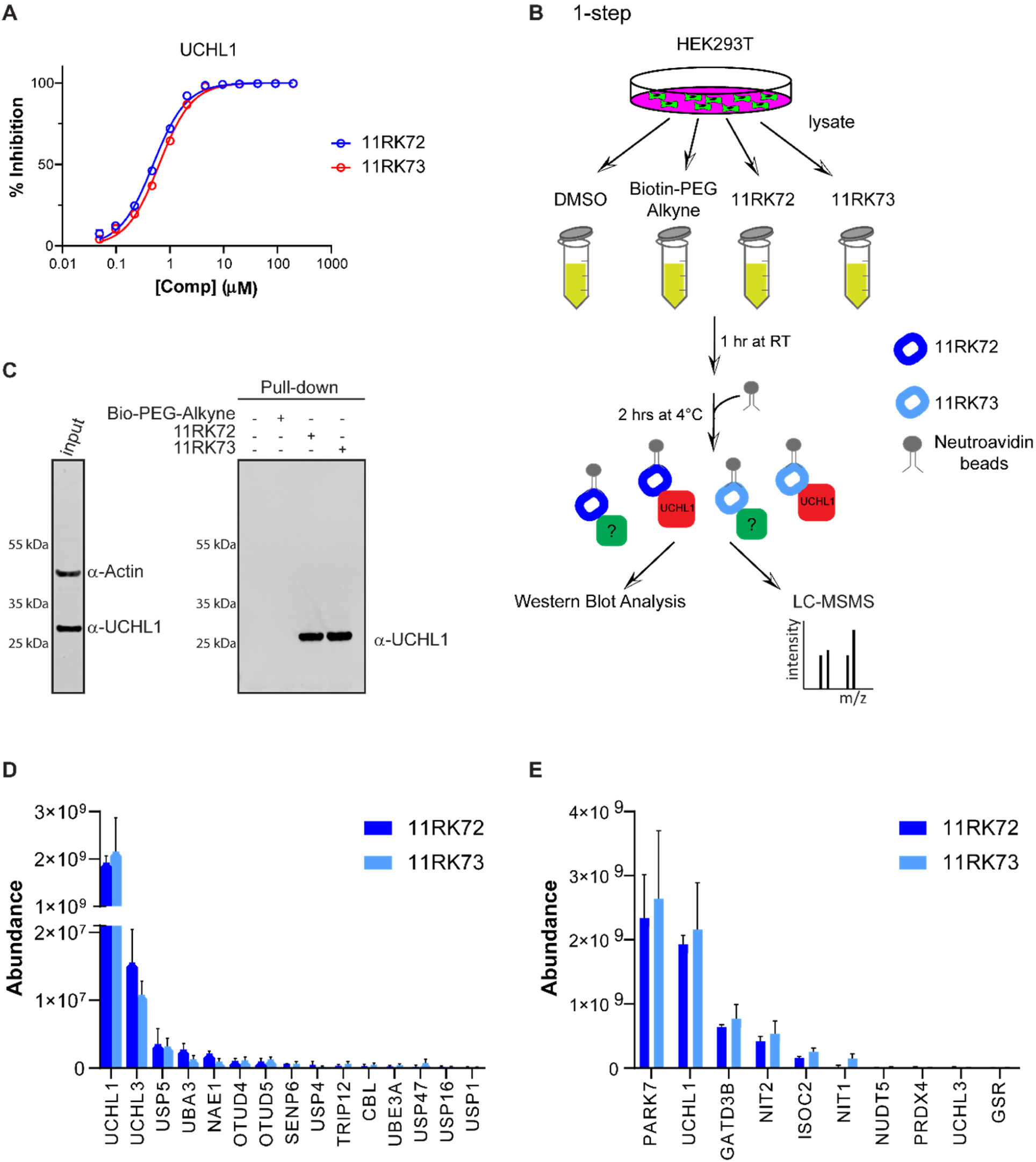
Proteomics experiments with biotinylated ABP analogs to identify ABP targets. A) IC_50_ determination of **11RK72** and **11RK73** for UCHL1. B) Schematic representation of pull-down experiment to identify ABP binding proteins. C) Confirmation of UCHL1 pull-down with biotinylated ABP analogs by Western Blot Analysis. Immunoblotting was performed using UCHL1 and Actin antibodies. Actin was used as a loading control and incubated together with UCHL1 antibody in the input sample. D) Abundances of the top-10 highest ranked proteins from the pull-down LC-MSMS experiment averaged over three replicates. E) Abundances of all enzymes related to the Ub(-like) system identified in the pull-down LC-MSMS experiment averaged over three replicates.

The abundances of the top-10 highest ranked proteins are shown in Figure 4E. In line with the results obtained with the 2-step approach, the highest ranked proteins are UCHL1 and PARK7. PARK7 shows a slightly higher abundance here, which contradicts our previous results from the in-cell labeling and 2-step pull-down experiments and might be attributed to the use of a different (biotinylated) version of the ABP or a peptide ionization difference during LC-MSMS measurements. The next highest ranked group of proteins, albeit at much lower abundance levels, includes two amidases NIT1 and NIT2, both harboring an active-site cysteine residue, the isochorismatase domain-containing protein 2 (ISOC2) and glutamine amidotransferase-like class-1 domain-containing protein 3B (GATD3B). Overall, the shorter (**11RK72**) and longer (**11RK73**) biotin probes give similar results, so the distance between probe and biotin does not seem to influence the binding nor the pull-down efficiency.

Upon comparison of the pull-down data (Figure 4) with the fluorescent probe labeling (Figure 3) we were unable to assign all bands to proteins. The majority of most abundant proteins in the pull-down experiment have a molecular weight between 20 and 35 kDa. Especially the pronounced band around 55 kDa in Figure 3D remains elusive. In a final attempt to assign this band we resolved the pull-down protein sample from the 1-step labeling experiment by SDS-PAGE. All proteins were visualized by silver staining after which the bands were excised and analyzed by LC-MSMS (Supporting Information Figure S2F). Again, UCHL1 and PARK7 were clearly the main proteins identified from the bands at ∼25 kDa. The proteins corresponding to the other bands were less clear but the main candidates were GAPDH at ∼40 kDa and Elongation factor 1α, tubulin or glutathione reductase (GSR) at ∼60 kDa. Whether or not these proteins actually bind to the probe or that these results are due to their high expression levels remains elusive. Based on the result that we identified UCHL1 as the major probe target in three individual experiments and that we found PARK7 as the only major off-target, we reasoned that **8RK59** could well be used for in-cell and *in vivo* labelling of UCHL1 activity.

### Probing UCHL1 activity in cells with 8RK59

To assess the application of **8RK59** in live cells, we used inverted fluorescent microscopy to image the **8RK59** signal in MDA-MB-436 and A549 cells after a 16-hour treatment with **8RK59**. Results showed that **8RK59** could penetrate and label the cells (Figure 5A). Compared to the control group, the BodipyFL signal was significantly decreased in UCHL1 knock-down MDA-MB-436 cells and similar results were observed for A549 cells (Supporting Information Figure S3). After imaging, MDA-MB-436 cells were lysed and followed with SDS-PAGE, fluorescence gel scanning and immunoblotting. A decreased UCHL1 signal was detected in MDA-MB-436 UCHL1 knock-down cells by **8RK59** and by antibody stain (Figure 5B). To further validate whether we can visualize UCHL1 specific activity inside cells, control and UCHL1 depleted A549 cells were pre-incubated with **8RK59** probe for 16 hours and stained with UCHL1 antibody (Figure 5C). We observed changes in the distribution of the probe inside the cells. In the control cells **8RK59** accumulated in both UCHL1-positive and negative subcellular compartments while in the UCHL1 knock-down cells the **8RK59** signal was largely decreased in the UCHL1-positive compartments, implying that UCHL1 binds to **8RK59** probe. In agreement with the gel–based labeling data shown in Figure 3 and the proteomics data shown in Figure 4 (and Supporting Information), we still observed some background subcellular localization of **8RK59** in UCHL1 knock-down cells, which may be the result of PARK7 staining. Taken all together, this result shows that the cellular distribution of **8RK59** probe changes upon depletion of UCHL1 and it can therefore be used to monitor UCHL1 activity in cells.

**Figure 5.**
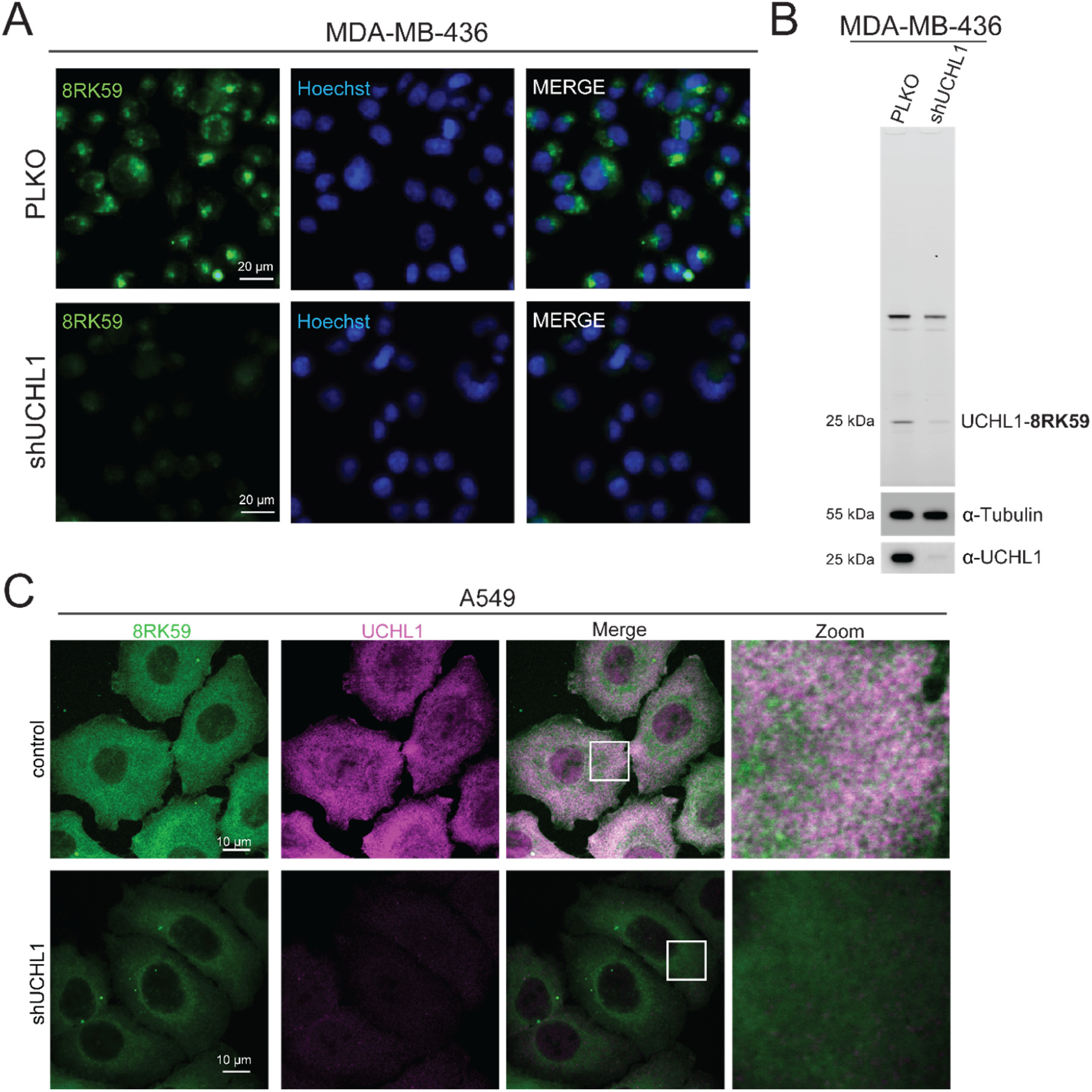
Probing UCHL1 activity in cells with **8RK59**. A) Live-cell fluorescence imaging of PLKO and shUCHL1 MDA-MB-436 cells treated with **8RK59**. B) Fluorescence labeling of endogenous UCHL1 in PLKO and shUCHL1 MDA-MB-436 cells treated with **8RK59** in SDS-PAGE gel. Immunoblotting was performed using UCHL1 antibody, and Tubulin was used as a loading control. C) Immunofluorescent staining of UCHL1 in **8RK59** pretreated PLKO and shUCHL1 A549 cells.

### Probing UCHL1 activity in zebrafish embryos with 8RK59

To investigate the application of **8RK59** in tracking UCHL1 activity in an *in vivo* model, we chose the zebrafish (*Danio rerio*) due to their high genetic homology to humans and the transparency of their embryos.^39^ Firstly, we treated zebrafish embryos with **8RK59** and recorded fluorescent images during the development of embryos from 1 to 7 days post fertilization (dpf). Results showed **8RK59** mainly labeled the nose, eye and brain of the zebrafish embryos (Figure 6A). Interestingly, all these organs are enriched in nerve cells and highly express *Uchl1* mRNA.^40^ To validate that the labelling of **8RK59** in zebrafish embryos is specific to UCHL1 protein, we fixed the **8RK59** labelled embryos and performed IF staining with UCHL1 antibody. Results demonstrated both the **8RK59** and UCHL1 antibody label similar organs of zebrafish embryos (Figure 6B). To assess whether **8RK59** could detect the UCHL1 activity changes in zebrafish embryos, we pretreated the zebrafish embryos with UCHL1 activity inhibitor **6RK73** from 1 to 3 dpf, and then labelled the embryos with **8RK59** from 4 to 6 dpf. We found that increasing concentrations of **6RK73**-pretreated zebrafish embryos resulted in significantly lower **8RK59** signal labelling (Figure 6C). In addition, the lysate of **6RK73**-pretreated zebrafish embryos showed decreased UCHL1 signal in fluorescence scans of a corresponding SDS-PAGE gel (Supporting Information Figure S4). These *in vivo* experiments indicate that **8RK59** can visualize and track UCHL1 activity during the development of zebrafish embryos.

**Figure 6.**
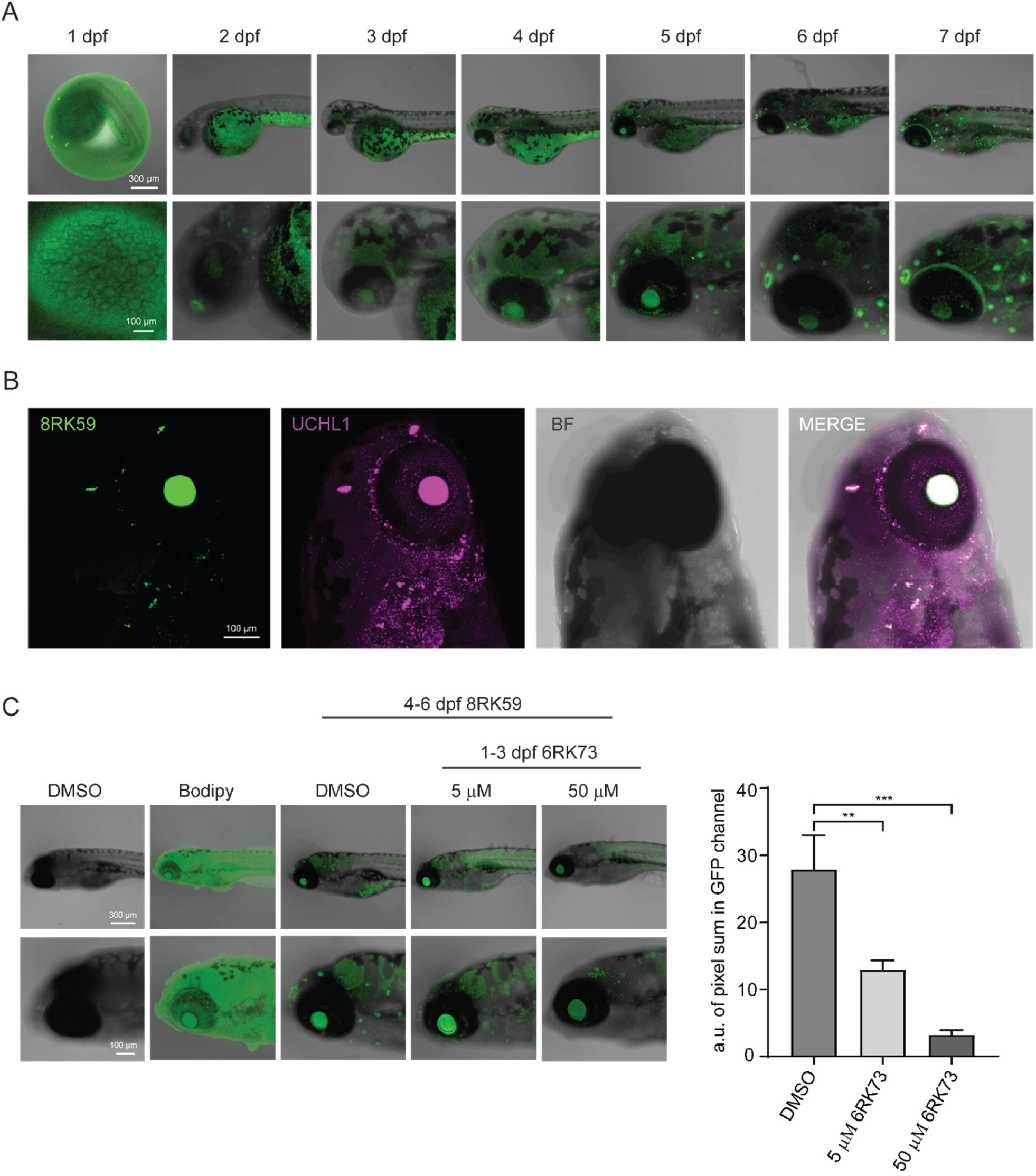
Probing UCHL1 activity in zebrafish embryos with **8RK59**. A) Tracking the localization of active UCHL1 with **8RK59** during the development of zebrafish embryos from 1 to 7 dpf. B) Immunofluorescent staining of UCHL1 in 6 dpf zebrafish embryo pretreated with **8RK59**. C) Monitoring UCHL1 activity changes in 6 dpf zebrafish embryos with **8RK59** pretreated with UCHL1 inhibitor **6RK73**. Representative images from five groups with zoom in of the brain area are shown in the left panel. The quantification of **8RK59** signal in three **6RK73** treatment groups are shown in the right panel. DMSO and BodipyFL dye were used as controls. **, P < 0.01, ***, P < 0.001, two-way ANOVA.

## CONCLUSIONS

One of the key challenges within DUB research is the creation of activity-based probes that target a single DUB type and at the same time are able to cross the cell membrane, in order to study these enzymes inside living cells or even living organisms.^41^ It has recently been shown by us and others that Ub-based tools (such as ABPs) can be made sub-type specific by engineering the amino acid sequence in Ub,^32, 42,43^ however these ABPs are not cell-permeable, although the use of cell-penetrating peptides has recently been applied to deliver Ub ABPs into cells.^44^ ABPs based on small-molecule inhibitors on the other hand are often cell-permeable and can be tuned chemically to become selective,^45, 46^ although such ABPs for DUBs have been lacking so far. We here provide evidence for the first fluorescent small-molecule target specific DUB ABP (**8RK59**) that hits UCHL1 activity *in vitro*, in cells and *in vivo*. We based our design on a cyanimide-containing inhibitor and show, in contrast to what has been reported in literature,^27^ that cyanimides can act as (near to) irreversible binders. Whether the irreversible bond formation results from the chemical nature of the cyanimide used here or from its binding mode within the UCHL1 active site and whether this property can be extended to other DUBs, remains to be investigated. Instalment of a fluorescent group onto a small-molecule inhibitor can have a detrimental effect on its inhibitory properties. Our data show that the installation of a Rhodamine fluorophore hardly, and a BodipyFL fluorophore only marginally effected the inhibitory potency towards UCHL1, whereas our Ub-ABP experiments confirmed the preservation of their selectivity for UCHL1 among other cysteine DUBs. From these two probes Rhodamine-tagged **9RK87** showed better *in vitro* characteristics, e.g. lower IC_50_ value and more potent in cell lysate, but unfortunately proved to be unable to cross the cell membrane. As such, this probe could be preferred for *in vitro* experiments and might be optimized for in-cell use by chemically improving the cell-penetrating properties of Rhodamine.^28^

Small-molecule inhibitors or probes almost inevitably result in unspecific interactors and this is not different for our compounds. We have considerably invested in the identification of potential off-targets of our probes by means of a proteomics approach. The data generated in this effort are not only useful for our own study but also provide valuable information for others working on this type of cyanimide-containing compounds. The proteomics data is in line with the Ub-probe experiments, confirming that these compounds are UCHL1 specific within the Ub system and to enzymes of the closely related Ub-like systems (e.g. Nedd8, SUMO, etc.). We indeed found a few potential off-targets, the main one being the protein and nucleotide deglycase PARK7. These cyanimide compounds may therefore provide a good starting point for small-molecule probes targeting PARK7, which, in spite of its important enzymatic function in protein and DNA repair in virtually any cell, have not been developed yet. Based on our data we expect that the potency and selectivity of the probe can be further improved by means of chemical alterations of the inhibitor. A better knowledge on the structural determinants of the interactions between probe and UCHL1 will be of great value for this, unfortunately despite several crystallization attempts we were unable to obtain appropriately diffracting crystals. During preparation of our manuscript Flaherty and co-workers^47^ reported on a related (*S*)-1-cyanopyrrolidine-2-carboxamide-based UCHL1 inhibitor and they applied NMR and molecular modeling to gain insight in the interactions between inhibitor and UCHL1, which could provide useful information to further optimize our probes. In addition, they modified their inhibitor with an alkyne moiety, which, unlike our molecules, resulted in a decrease in potency towards UCHL1 and selectivity with respect to UCHL3. This 2-step probe was then used to identify off-targets in KMS11 cells but remarkably none of their identified proteins show overlap with our list.

In conclusion, we have developed a fluorescent small-molecule activity-based probe that labels UCHL1 activity *in vitro*, in cells and *in vivo*. It is the first example of a ‘1-step’ DUB-selective, cell-permeable ABP and therefore serves as a unique addition to the ‘Ub toolbox’, concomitantly addressing two of the outstanding challenges within this field. Our results show that the probe works in several different cell lines and we therefore foresee a potential wide application of the probe in studying UCHL1 activity related to neurodegenerative disorders and cancer. In fact, we recently showed that **6RK73** decreases UCHL1 activity and thereby inhibits TGFβ/SMAD2 and SMAD3 signaling and breast cancer migration and extravasation.^48^ We are convinced that the here reported strategy of small-molecule cyanimide-based probes can be expanded to other cysteine proteases and specifically DUBs. With the rising importance of the Ub system as source of practical drug targets we believe that these ABP tools will fill an unmet need allowing us to study active DUBs in their native environment in live cells or animals and as such aid in the development of future therapeutics that target diseases associated with ubiquitination.

## METHODS

### IC_50_ determination

The *in vitro* enzyme inhibition assays were performed in “non-binding surface flat bottom low flange” black 384-well plates (Corning) at room temperature in a buffer containing 50 mM Tris·HCl, 100 mM NaCl, pH 7.6, 2.0 mM cysteine, 1 mg/mL 3-[(3-cholamidopropyl) dimethylammonio] propanesulfonic acid (CHAPS) and 0.5 mg/mL γ-globulins from bovine blood (BGG) in triplicate. Each well had a final volume of 20.4 µL. All dispensing steps involving buffered solutions were performed on a Biotek MultiFlowFX dispenser. The compounds were dissolved in DMSO as 10 mM, 1 mM and 0.1 mM stock solutions and appropriate volumes were transferred from these stocks to the empty plate using a Labcyte Echo550 acoustic dispenser and accompanying dose-response software to obtain a 12 point serial dilution (3 replicates) of 0.05 to 200 µM. A DMSO back-fill was performed to obtain equal volumes of DMSO (400 µL) in each well. 10 mM *N*-ethylmaleimide (NEM) was used a positive control (100% inhibition) and DMSO as negative control (0% inhibition). 10 µL buffer was added and the plate was vigorously shaken for 20 sec. Next, 5 µL of a 4× final concentration enzymes stock was added followed by incubation for 30 min. 5 µL of the substrate (Ub-Rho-morpholine (final concentration 400 nM) or Cbz-PheArg-AMC (final concentration 10 µM) in the case of Papain) and the increase in fluorescence intensity over time was recorded using a BMG Labtech CLARIOstar or PHERAstar plate reader (excitation 487 nm, emission 535 nm). The initial enzyme velocities were calculated from the slopes, normalized to the positive and negative controls and plotted against the inhibitor concentrations (in M) using the built-in equation “[inhibitor] vs. response – Variable slope (four parameters), least squares fit” with constraints “Bottom = 0” and “Top = 100” in GraphPad Prism 7 software to obtain the IC_50_ values.

### Jump dilution assay

All assays were performed in triplicate. The assay was performed in a buffer containing 50 mM Tris·HCl, 100 mM NaCl, pH 7.6, 2.0 mM cysteine, 1 mg/mL 3-[(3-cholamidopropyl) dimethylammonio] propanesulfonic acid (CHAPS) and 0.5 mg/mL γ-globulins from bovine blood (BGG). The final concentrations used were: 3 nM UCHL1, 400 nM Ub-Rho-morpholine, 10 µM or 0.1 µM or a jump dilution of 10 µM to 0.1 µM inhibitor. Samples of 20 µL containing 300 nM UCHL1 and 20 µM inhibitor (2% DMSO), 2% DMSO or 20 mM *N*-ethylmaleimide (NEM) were incubated for 30 min. at room temperature. 5 µL of each sample was then diluted into a 500 µL solution containing 400 nM Ub-Rho-morpholine. After a brief mixing 20 µL of each of these solutions was quickly transferred to a “non-binding surface flat bottom low flange” black 384-well plate (Corning) and the increase in fluorescence over time was recorded using a BMG Labtech CLARIOstar plate reader (excitation 487 nm, emission 535 nm). As a control, samples were taken along in which 40 µL of a 20 µM and 0.2 µM inhibitor solution in buffer (2% DMSO) were added to 20 µL of a 12 nM UCHL1 solution. After 30 min. incubation 20 µL of a 1.6 µM Ub-Rho-morpholine solution was added after which 20 µL of each solution was transferred to the same 384 well plate mentioned above and the increase in fluorescent intensity was measured concomitantly. Fluorescent intensities were plotted against time using GraphPad Prism 7.

### Covalent complex formation mass spectrometry analysis

Samples of 1.4 µM UCHL1 in 70 µL buffer containing 50 mM Tris·HCl, 100 mM NaCl, pH 7.6, 2.0 mM cysteine and 1 mg/mL 3-[(3-cholamidopropyl) dimethylammonio] propanesulfonic acid (CHAPS) were prepared. These samples were treated with 1 µL DMSO or 1 µL of a 10 mM inhibitor/probe stock solution in DMSO (140 µM final concentration) and incubated for 30 min. at room temperature. Samples were then 3× diluted with water and analyzed by mass spectrometry by injecting 1 µL on a Waters XEVO-G2 XS Q-TOF mass spectrometer equipped with an electrospray ion source in positive mode (capillary voltage 1.2 kV, desolvation gas flow 900 L/hour, T = 60 ^o^C) with a resolution R = 26,000. Samples were run using 2 mobile phases: A = 0.1% formic acid in water and B = 0.1% formic acid in CH_3_CN on a Waters Acquity UPLC Protein BEH C4 column, 300 Å, 1.7 µm (2.1 × 50 mm); flow rate = 0.5 mL/min, runtime = 14.00 min, column T = 60 °C, mass detection 200-2500 Da. Gradient: 2 – 100% B. Data processing was performed using Waters MassLynx Mass Spectrometry Software 4.1 and ion peaks were deconvoluted using the built-in MaxEnt1 function.

### Probe labeling of purified recombinant UCHL1

The assay was performed in a buffer containing 50 mM Tris·HCl, 100 mM NaCl, pH 7.6, 2.0 mM cysteine and 1 mg/mL 3-[(3-cholamidopropyl) dimethylammonio] propanesulfonic acid (CHAPS). A stock solution containing 8 µM UCHL1 and stock solutions containing 20 µM **8RK59, 9RK15, 9RK87** and Rho-Ub-PA in buffer were prepared. 50 µL of the UCHL1 stock solution was mixed with 50 µL of all probe solutions followed by incubation for 60 min. at 37 °C. Three aliquots of 10 µL of each sample were taken and treated with 1) 5 µL loading buffer with β-mercaptoethanol, followed by 5 min. heating at 95 °C; 2) 5 µL loading buffer with 50 mM TCEP; 3) 5 µL loading buffer. Samples were resolved by SDS-PAGE using a 4-12% Bis-Tris gel (Invitrogen, NuPAGE) with MES SDS running buffer (Novex, NuPAGE) for 45 min. at 190V. Gels were scanned for fluorescence on a GE Typhoon FLA 9500 using a green (λ_ex/em_ 473/530 nm) and red (λ_ex/em_ 532/570 nm) channel followed by staining with InstantBlue Coomassie protein stain (Expedeon) after which the gel was scanned on a GE Amersham Imager 600.

### Cell lines and cell culture

HEK293T, HeLa, A549 and MDA-MB-436 cells were originally obtained from American Type Culture Collection (ATCC) and SKBR7 cells were obtained from Dr. J. Martens (Erasmus University Medical Center, Rotterdam, The Netherlands). Cells were cultured in Dulbecco’s modified Eagles’s medium (DMEM) supplemented with 10% fetal bovine serum (FBS) and 100 U/mL penicillin-streptomycin (15140122; Gibco). Stable shUCHL1 A549 and shUCHL1 MDA-MB-436 cell lines were generated by lentiviral infection and the cell lines were continuously cultured under puromycin selection. Four UCHL1 shRNAs were identified and tested, the most effective shRNA (TRCN0000007273; Sigma) for lentiviral infection were used for experiments. All cell lines were regularly tested for absence of mycoplasma and were authenticated.

### Transfection

For shRNA expression, lentiviruses were produced by transfecting shRNA-targeting plasmids together with helper plasmids pCMV–VSVG, pMDLg–RRE (gag–pol), and pRSV–REV into HEK293T cells. Cell supernatants were collected 48 hours after transfection and were used to infect cells. To obtain stable shUCHL1 A549 and shUCHL1 MDA-MB-436 UCHL1 knock-down cell lines, cells were infected at low confluence (40%) for 12 hours with lentiviruses in the presence of 5 ng/mL Polybrene (Sigma). Cells were subjected to 1 μg/mL puromycin selection 48 hours after infection. Four UCHL1 shRNAs were identified and tested, the most effective UCHL1 shRNA (TRCN0000007273; Sigma) for lentiviral infection was used for the experiments.

For siRNA transfection, siRNAs targeting UCHL1 (Set of 4: siGENOME; MQ-004309-00-0002 2 nmol) and PARK7 (Set of 4: siGENOME; MQ-005984-00-0002 2 nmol) were obtained from Dharmacon. Knock-down of UCHL1 and PARK7 in HEK293T cells was performed as follows: for 6-well plate format 200 µL siRNA (500 nM stock) were incubated with 4 µL Dharmafectin reagent 1 (Dharmacon) diluted in 200 µL medium without supplements by shaking for 20 min. at room temperature. The transfection mix was added to cells and cultured at 37 °C and 5 % CO_2_. 48 hours after transfection **8RK59** was added to the cells and incubated for 24 hours. Cells were harvested and analysed as described under the section “DUB activity profiling and competition with Ub-PA DUB probes”.

For the expression of UCHL1 in HEK293T cells, Flag-HA-UCHL1 construct was obtained from Addgene (22563). Catalytically inactive mutant (C90A) UCHL1 was generated using site-directed mutagenesis. Wild-type and C90A mutant UCHL1 were transfected into HEK293T cells using PEI transfection reagent. 24 hours after transfection **8RK59** was added to the cells and incubated for 24 hours. Cells were harvested and analysed as described under the section “DUB activity profiling and competition with Ub-PA DUB probes”.

### Immunoblotting

Cells were lysed in HR lysis buffer (50 mM Tris, 5 mM MgCl_2_, 250 mM sucrose and 2 mM DTT, pH 7.4) with protease inhibitor cocktail for 10 min. on ice. The lysates were sonicated using 10 cycles of 30 sec. pulse on, 30 sec. pulse off. The lysates were centrifuged at maximun speed for 20 min. at 4 °C, thereafter protein concentrations were measured using the DC protein assay (500-0111; Bio-Rad) and equal amounts of proteins were used for each condition that was analyzed by immunoblotting with following antibodies: UCHL1 (ab27053; Abcam), Tubulin (2148; Cell Signaling), GAPDH (MAB374; Millipore), Actin (A5441; Sigma-Aldrich,).

### Immunofluorescence staining

Cells were fixed for 20 min. in 4% paraformaldehyde and then permeabilized in 0.1% Triton-X for 10 min. Non-specific binding was blocked with blocking buffer (1% BSA in 0.1% PBS-Tween) for 30 min. The primary antibody UCHL1 (ab27053; Abcam) was diluted in blocking buffer and added to the cell for 1 hour. After 3 times washing with PBS, the secondary antibody donkey anti rabbit IgG Alexa Fluorescence 555 (Invitrogen #A31572) was added and incubated for 30 min. After 3 times washing with PBS, samples were mounted with VECTASHIELD antifade mounting medium with DAPI (H-1200; Vector Laboratories). Fluorescence images were acquired with TCS SP8 confocal microscope (Leica). Zebrafish embryos were fixed with 4% paraformaldehyde 2 hours at room temperature. Samples were dehydrated with 33%, 66%, 100% methanol in PBS, followed by a rehydration step. Thereafter, the embryos were successively treated with 10 μg/mL proteinase K for 60 min. at 37 °C, permeabilized with 0.25% Triton in PBS for 30 min. on ice, and blocked with 10% FBS in PBS for 1 hour at room temperature. Embryos were incubated with primary antibody (ab27053; Abcam) for at least 12 hours at 4 °C. After washing with 0.1% Triton in PBS for 3 times 10 min., the samples were incubated with fluorescein-conjugated secondary antibody donkey anti rabbit IgG Alexa Fluorescence 555 (Invitrogen #A31572) for 2 hours at room temperature. After washing with PBS (0.1% Triton), samples were analyzed using a confocal microscope SP5 STED (Leica, Rijswijk, Netherlands).

### DUB activity profiling and competition with Ub-PA DUB probes

HEK293T cells were treated with 5 µM final concentration of the indicated compounds for 24 hours. Cells were lysed in HR lysis buffer supplemented with protease inhibitor cocktail (11836145001; Roche). Samples were kept on ice and lysed by sonication (10 cycles of 30 seconds on and 30 seconds off). 25 μg protein extract was labelled with either 1 μM Rh-Ub-PA probe or 0.5 μM Cy5-Ub-PA probe for 30 min. at 37 °C. For the cell lysate incubation, HEK293T cells were lysate as described above. HEK293T cell lysates were preincubated with 5 µM final concentration of compounds for 1 hour, followed by incubation with 0.5 μM Cy5-Ub-PA probe for 30 min. at 37 °C. Labelling reactions were terminated with sample buffer and heating to 100 °C for 10 min. Samples were size-separated in SDS-PAGE gels. In-gel fluorescence signals were scanned employing the Typhoon FLA 9500 Molecular Imager (GE Healthcare). Images were analyzed using ImageJ software.

### Probe labelling of endogenous UCHL1 in living cell

Cell lines were transfected with shRNAs, siRNAs or UCHL1 constructs as described above. 5 µM final concentration of probes were added to the cell a day before harvesting. Cells were harvested in HR buffer as described above. NuPAGE LDS sample buffer containing 50 mM TCEP was added to cell lysates. Samples were resolved by SDS-PAGE using a 4-12% Bis-Tris gel (Invitrogen, NuPAGE) with MES SDS running buffer (Novex, NuPAGE) for 45 min. at 190V. Gels were scanned for fluorescence on a GE Typhoon FLA 9500 using a green (λ_ex/em_ 473/530 nm) and red (λ_ex/em_ 532/570 nm) channel followed by transferring proteins to nitrocellulose membrane (Amersham) and western blot analysis.

### Proteomics

For 1-step approach, 4 × 10^6^ HEK293T cells were seeded into 10 cm dishes for each treatment. 48 hours later, HEK293T cells were harvested in lysis buffer containing 50 mM HEPES pH 7.3, 150 mM NaCl and 1% NP-40 and 1× protease inhibitor cocktail and incubated for 30 min. on ice. Cell lysates were centrifuged at maximum speed for 20 min. The lysates were incubated with 5 µM final concentration of Biotin-PEG_4_-alkyne, **11RK72** or **11RK73** or same volume of DMSO for 1 hour at room temperature. 30 µL of neutravidin beads slurry (50%) were added to each sample. The samples were then incubated for 2 hours at 4 °C. Beads were washed six times in wash buffer containing 50 mM HEPES pH 7.3, 150 mM NaCl and 1% NP-40. After completely removing the washing buffer, NuPAGE LDS sample buffer (containing 7.5% β-mercaptoethanol) was added to the beads followed by 15 min. incubation at 95 °C.

For 2-step approach, 4 × 10^6^ HEK293T cells were seeded into 10 cm dishes for each treatment. 24 hours later, 5 µM final concentration of **8RK64** or same volume of DMSO was added to the cells. After 24 hours incubation, HEK293T cells were harvested in lysis buffer containing 50 mM HEPES pH 7.3, 150 mM NaCl and 1% NP-40 and 1× protease inhibitor cocktail and incubated for 30 min. on ice. Cell lysates were centrifuged at maximum speed for 20 min. 1× volume of click cocktail (100 mM CuSO_4._5H_2_O, 1M sodium ascorbate, 100 mM TBTA (Tris[(1-benzyl-1*H*-1,2,3-triazol-4-yl)methyl]amine) ligand, 0.1 M HEPES pH 7.3 and 5 µM biotin-alkyne) were added to 2× volume of cell lysates and incubated for 45 min. 30 µL of neutravidin beads slurry (50%) were added to each sample. The samples were then incubated for 2 hours at 4 °C. Beads were washed six times in wash buffer containing 50 mM HEPES pH 7.3, 150 mM NaCl and 1% NP-40. After completely removing the washing buffer, SDS sample buffer (containing 7.5% β-mercaptoethanol) was added to the beads followed by 15 min. incubation at 95 °C. For MS analysis proteins were run for 1-2 cm on a 4-12% PAGE (NuPAGE Bis-Tris Precast Gel, Life Technologies) and stained with silver (SilverQuest Silver Stain, Life Technologies). The lane was cut into two equal parts, and gel slices subjected to reduction with dithiothreitol, alkylation with iodoacetamide and in-gel trypsin digestion using a Proteineer DP digestion robot (Bruker).

Tryptic peptides were extracted from the gel slices, lyophilized, dissolved in 95/3/0.1 v/v/v water/acetonitril/formic acid and subsequently analyzed by on-line C18 nanoHPLC MS/MS with a system consisting of an Easy nLC 1000 gradient HPLC system (Thermo, Bremen, Germany), and a LUMOS mass spectrometer (Thermo). Fractions were injected onto a homemade precolumn (100 μm × 15 mm; Reprosil-Pur C18-AQ 3 μm, Dr. Maisch, Ammerbuch, Germany) and eluted via a homemade analytical nano-HPLC column (15 cm × 50 μm; Reprosil-Pur C18-AQ 3 µm). The gradient was run from 0% to 50% solvent B (20/80/0.1 water/acetonitrile/formic acid v/v/v) in 20 min. The nano-HPLC column was drawn to a tip of ∼5 μm and acted as the electrospray needle of the MS source. The LUMOS mass spectrometer was operated in data-dependent MS/MS (top-10 mode) with collision energy at 32 V and recording of the MS2 spectrum in the orbitrap. In the master scan (MS1) the resolution was 120,000, the scan range 400-1500, at an AGC target of 400,000 @maximum fill time of 50 ms. Dynamic exclusion after n=1 with exclusion duration of 10 s. Charge states 2-5 were included. For MS2 precursors were isolated with the quadrupole with an isolation width of 1.2 Da. HCD collision energy was set to 32V. First mass was set to 110 Da. The MS2 scan resolution was 30,000 with an AGC target of 50,000 @maximum fill time of 60 ms.

In a post-analysis process, raw data were first converted to peak lists using Proteome Discoverer version 2.2.0.388 (Thermo Electron), and then submitted to the Uniprot Homo sapiens database (67911 entries), using Mascot v. 2.2.04 (www.matrixscience.com) for protein identification. Mascot searches were with 10 ppm and 0.02 Da deviation for precursor and fragment mass, respectively, and trypsin as enzyme. Up to two missed cleavages were allowed, and methionine oxidation was set as a variable modification; carbamidomethyl on Cys was set as a fixed modification. Protein abundance calculation and statistical analysis was performed using Proteome Discoverer software.

### Zebrafish experiments

Transgenic zebrafish lines Tg (kdrl: mTurquois) were raised, staged and maintained according to standard procedures in compliance with the local Institutional Committee for Animal Welfare of the Leiden University. Zebrafish embryos were treated with 5 µM **8RK59** or gradient **6RK73** concentration in the egg water. Fluorescent image acquisition was performed with a Leica SP5 STED confocal microscope (Leica, Rijswijk, Netherlands). The quantification of **8RK59** signal was analyzed by Leica microscope software platform LAS X. 30 zebrafish were treated in each group and 3 representative images were taken and analyzed. Statistical analysis was performed using Graphpad Prism 8 software. Numerical data from triplicates are presented as the mean ± SD. Two-way analysis of variance (ANOVA) has been used to analyze multiple subjects.

## Supporting information

The list of identified proteins

## ASSOCIATED CONTENT

### Supporting Information

The Supporting Information is available free of charge DOI: ### Detailed synthesis procedures and compound characterization (including Ub-Rho-morpholine and Rhodamine110-alkyne), NMR and LC-MS spectral data, proteomics data, IC_50_ data and supporting information figures.

## AUTHOR INFORMATION

### Notes

The authors declare the following competing financial interest(s): H.O. is shareholder of the reagent company UbiQ.

### Ethics Statement

Zebrafish were raised, staged and maintained according to standard procedures in compliance with the local Institutional Committee for Animal Welfare of the Leiden University.

## ACKNOWLEDGEMENTS

We thank Yves Leestemaker for assistance with the DUB activity-based profiling assays and Bjorn van Doodewaerd for LC-MS and HPLC purification assistance. P.t.D. and H.O. are supported by Oncode Institute. S.L. is supported by the China Scholarship Council. This work was supported by the Dutch Organization for Scientific Research NWO VICI grant (724.013.002) to H.O., Investment Grant NWO Medium grant (91116004, partly financed by ZonMw) to P.A.v.V. and Cancer Genomics Centre Netherlands (CGC.NL) to P.t.D.

## FIGURE LEGENDS

**Scheme 1.**
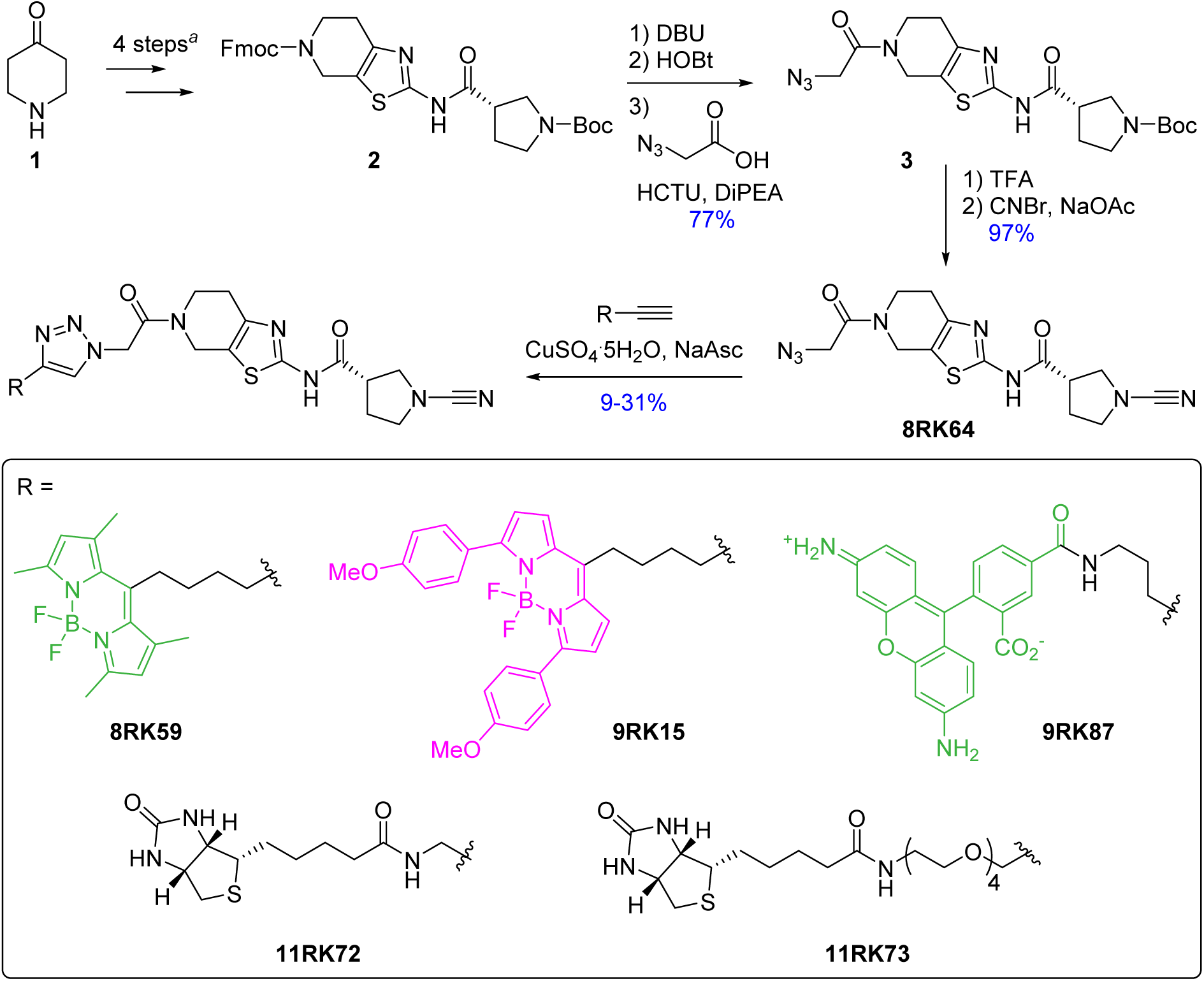
Synthesis of azide-containing UCHL1 inhibitor 8RK64 and fluorescent and biotinylated probe derivatives thereof. ^*a*^ Synthetic steps described in literature.^26^

