## Supporting Information for "A Small-Molecule Activity-Based Probe for Monitoring Ubiquitin C-terminal Hydrolase L1 (UCHL1) Activity in Live Cells and Zebrafish Embryos"

**Contents**

Table S1. $IC_{50}$ values 2

Figure S1. Cell permeability of **8RK59**, **9RK15**, and **9RK87** probes 3

Figure S2. Pull-down and proteomics analysis 4

Figure S3. Probing UCHL1 activity in cells with **8RK59** 6

Figure S4. Endogenous UCHL1 labeling in zebrafish embryo lysates 7

Compound synthesis 8

References 19

MS analysis of UCHL1 – compound binding 20

$IC_{50}$ curves 22

NMR & LC-MS analysis of synthesized compounds 23

**Table S1: $IC_{50}$ values[a]**

| Compound | DUB (concentration) | $IC_{50}$ (μM) |
|---|---|---|
| 6RK73 | UCHL1 (1 nM) | 0.23 |
| 6RK73 | UCHL3 (0.01 nM) | 235 |
| 6RK73 | UCHL5 (1 nM) | >>100 |
| 6RK73 | USP7 (1 nM) | 68.8 |
| 6RK73 | USP16 (2 nM) | >>100 |
| 6RK73 | USP30 (10 nM) | 9.8 |
| 6RK73 | Papain (3 nM) | 10.7 |
| 8RK64 | UCHL1 (1 nM) | 0.32 |
| 8RK64 | UCHL3 (0.01 nM) | 216 |
| 8RK64 | UCHL5 (1 nM) | >>100 |
| 8RK59 | UCHL1 (1 nM) | 1.2 |
| 9RK15 | UCHL1 (1 nM) | 3.6 |
| 9RK87 | UCHL1 (1 nM) | 0.44 |
| 11RK72 | UCHL1 (1 nM) | 0.50 |
| 11RK73 | UCHL1 (1 nM) | 0.64 |

[a] After 30 min. of incubation of enzyme with inhibitor.

### Figure S1

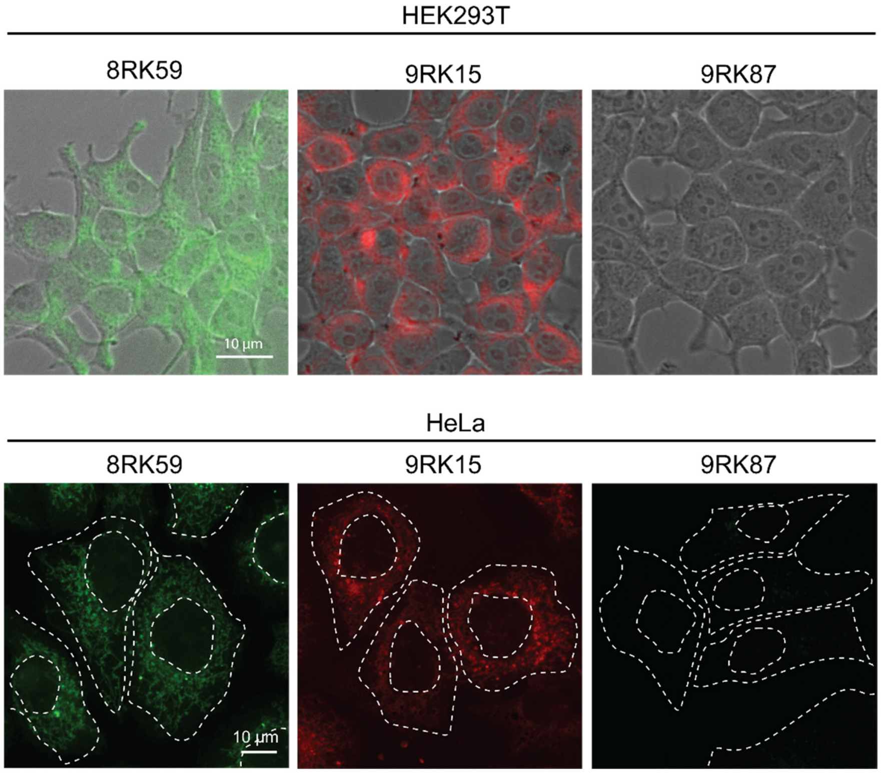


**Figure S1.** Cell permeability of **8RK59**, **9RK15**, and **9RK87** probes. HEK293T (top panel) and HeLa (bottom panel) cells were incubated with 5 µM final concentration of indicated probes for 24 hours at 37 °C. HEK293T cells were visualized using EVOS® FL Cell Imaging System. Nikon Plan Fluor 40×/0.75, infinity/0.17 objective was used. HeLa cells were fixed by 3.7% formaldehyde and then mounted using ProLong Gold antifade mounting medium with DAPI. Nuclei and cell boundaries are shown in dashed white lines. HeLa cells were imaged using Leica SP8 microscopes. HCX PL 63× 1.32 oil objectives and HyD detectors were used in confocal images.

### Figure S2

A

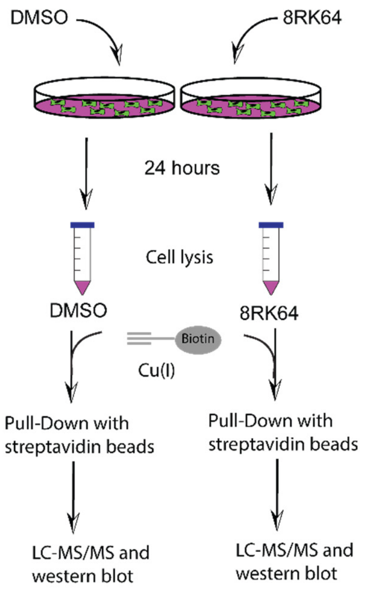


B

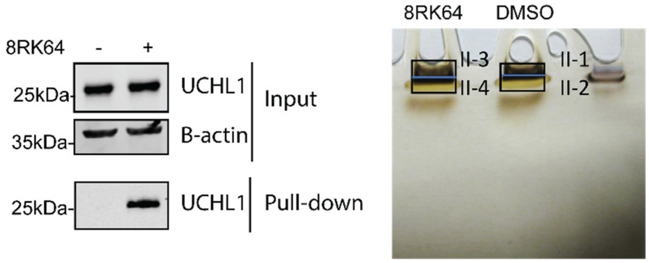


C

| | Identified Proteins | MW | DMSO | 8RK64 |
|---|---|---|---|---|
| 1 | Ubiquitin carboxyl-terminal hydrolase isozyme L1 OS=Homo sapiens | 25 kDa | | 15 |
| 2 | Actin, cytoplasmic 2 OS=Homo sapiens GN=ACTG1 | 42 kDa | 17 | 24 |
| 3 | Protein DJ-1 OS=Homo sapiens GN=PARK7 | 20 kDa | | 4 |
| 4 | Alpha-enolase OS=Homo sapiens GN=ENO1 | 47 kDa | 18 | 8 |
| 5 | Protease serine 1 OS=Homo sapiens GN=PRSS1 | 25 kDa | 11 | 11 |
| 6 | Tubulin alpha-1B chain OS=Homo sapiens GN=TUBA1B | 50 kDa | 9 | 12 |
| 7 | Elongation factor 1-alpha 1 OS=Homo sapiens GN=EEF1A1 | 50 kDa | 11 | 10 |
| 8 | Elongation factor 2 OS=Homo sapiens GN=EEF2 | 95 kDa | 8 | 12 |

D

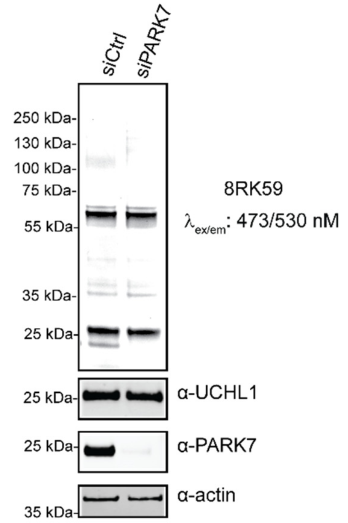


E

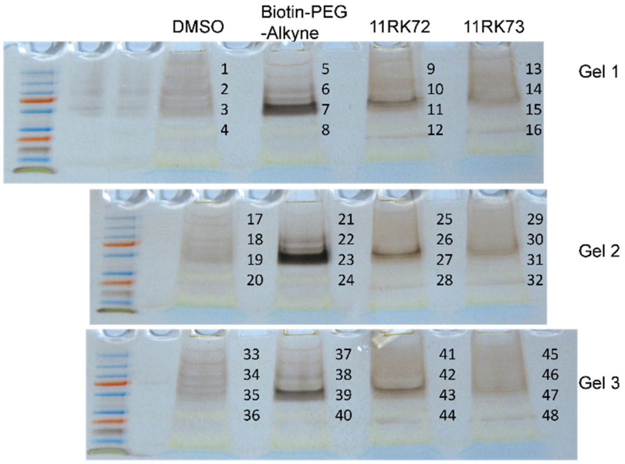


F

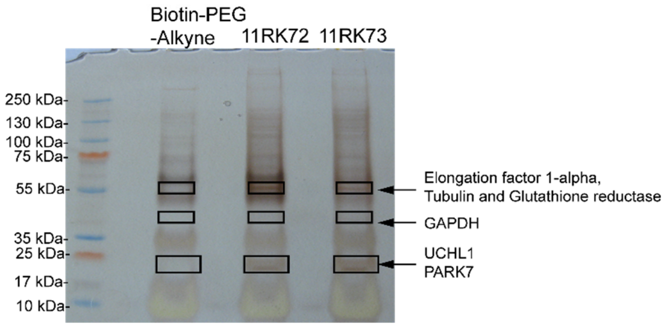

**Figure S2.** Pull-down and proteomics analysis either via a 1-step approach with biotin tagged-probes or via a 2-step labeling approach with click-chemistry. A) Schematic representation of 2-step labeling and pull-down approach. B) Western blot (left panel) and silver staining (right panel) of the samples obtained from 2-step labeling approach. Neutravidin beads from DMSO or **8RK64** treated samples were boiled in NuPAGE LDS sample buffer at 95 °C for 15 min. Proteins were run on 4-12% SDS-PAGE (1 cm for right panel) and either immunoblotted against PARK7, UCHL1 and Actin or stained using SilverQuest Silver Stain. The numbers on the gel indicate slices cut from the gel. C) List of the top eight proteins identified from proteomics experiment using 2-step labeling and pull-down approach. D) Confirmation of PARK7 labeling with **8RK59** via Fluorescent labeling and western blot analysis. Labeled proteins with **8RK59** were analyzed using in-gel fluorescence scanning followed by immunoblot against UCHL1. Actin is used as a loading control. E) Silver staining of three independent gels obtained from 1-step labeling approach. Related to Figure 4. Neutravidin beads from DMSO, Biotin-PEG$_4$-Alkyne, **11RK72** or **11RK73** treated samples were boiled in NuPAGE LDS sample buffer at 95 °C for 15 min. Proteins were run for 2 cm on 4-12% SDS-PAGE and stained using SilverQuest Silver Stain. The numbers on the gels indicate slices cut from the gels. F) Silver staining of the samples obtained from 1-step labeling approach**.** Neutravidin beads from Biotin-PEG$_4$-Alkyne, **11RK72** or **11RK73** treated samples were boiled in NuPAGE LDS sample buffer at 95 °C for 15 min. Proteins were run on 4-12% SDS-PAGE and stained using SilverQuest Silver Stain. Rectangles indicate the gel slices analyzed by LC-MS/MS.

### Figure S3

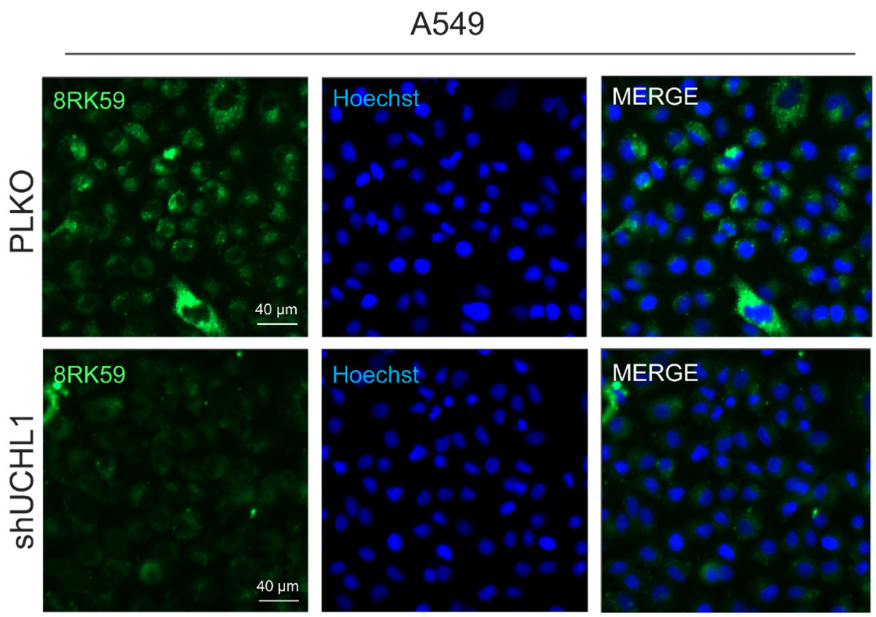


**Figure S3.** Probing UCHL1 activity in cells with **8RK59**. A549 PLKO and shUCHL1 cells were incubated with 5 µM **8RK59** for 16 hours at 37 °C. 1 µg/mL Hoechst 33342 was used to stain the nuclei of the live cells for 30 min. at 37 °C. Images were acquired with Leica DMi8 Inverted Fluorescent Microscope. HC PL FLUOTAR L 40x/0.60 DRY was used.

### Figure S4

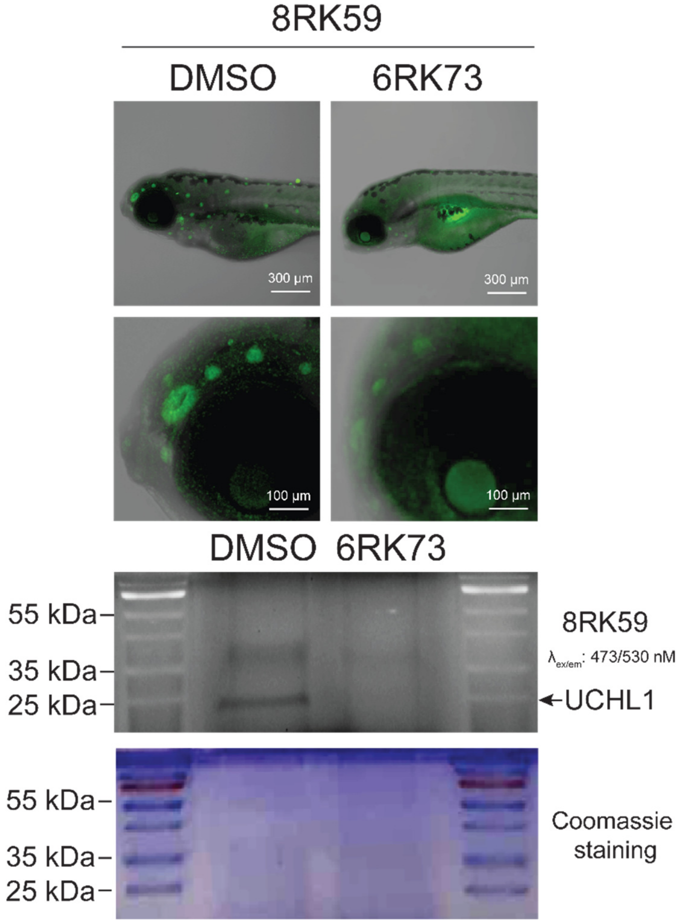


**Figure S4.** Fluorescence labeling of endogenous UCHL1 in zebrafish embryo and lysates with **8RK59**. Images of **8RK59** signal in zebrafish embryos pre-treated with DMSO or 5 µM **6RK73** (top panel). Fluorescence scan of corresponding zebrafish lysate in SDS-PAGE gel (bottom panel). Coomassie staining serves as loading control.

#### Compound synthesis

##### General

General reagents were purchased from Sigma-Aldrich, Biosolve, and Acros, and used as received. Solvents were purchased from Biosolve or Sigma-Aldrich. (*S*)-1-Boc-pyrrolidine-3-carboxylic acid (CAS number 140148-70-5, article number OR-5566) was purchased from Combi-Blocks Inc. BiotinPEG$_4$-alkyne (CAS number 1262681-31-1, article number PEG4950) was purchased from Iris Biotech GmbH). Thin Layer Chromatography (TLC) was performed on Merck aluminum sheets (pre-coated with silica gel 60 $F_{254}$). Compounds were visualized by UV adsorption (254 nm) and by using a solution of $KMnO_4$ (7.5 g $L^{-1}$) and $K_2CO_3$ (50 g $L^{-1}$) in $H_2O$ or a solution of ninhydrin (15 g $L^{-1}$) in 3% AcOH/EtOH v/v. Compounds (unless stated otherwise) were purified by a Büchi Sepacore automatic flash chromatography system X10/X50. The Büchi Sepacore system was equipped with two Büchi pump modules C-605, a Büchi control unit C-620, Büchi fraction collector C-660 and a Büchi UV Photometer C-640. The silica columns were purchased at GraceResolv™ and were packed with a grade of Davisil® silica. NMR spectra ($^{1}H$, $^{13}C$) were recorded on a Bruker Ultrashield 300 MHz spectrometer at 298 K. Resonances are indicated with symbols 'd' (doublet), 's' (singlet), 't' (triplet) and 'm' (multiplet). Chemical shifts (δ) are given in ppm relative to $CDCl_3$, DMSO-$d_6$ or $CD_3OD$ as an internal standard and coupling constants (*J*) are quoted in hertz (Hz). LC-MS measurements were performed on an LC-MS system equipped with a Waters 2795 Separation Module (Alliance HT), a Waters 2996 Photodiode Array Detector (190–750 nm), an Xbridge C18 column (2.1 × 100 mm, 3.5 μm) and an LCT ESI-Orthogonal Acceleration Time of Flight Mass Spectrometer. Samples were run using 2 mobile phases: A = 1% $CH_3CN$ and 0.1% formic acid in $H_2O$ and B = 1% $H_2O$ and 0.1% formic acid in $CH_3CN$. Data processing was performed using Waters MassLynx Mass Spectrometry Software 4.1. LC-MS Program: Waters Xbridge C18 column (2.1 × 100 mm, 3.5 μm); flow rate = 0.4 mL $min^{-1}$, runtime = 13 min, column T = 40 °C, mass detection: 100–1500 Da. Gradient: 0–0.4 min: 5% B; 0.4–9.0 min: 5% → 95% B; 9.0–11.2 min: 95% B; 11.2–11.3 min: 95% → 5% B; 11.3–13.00 min: 5% B. Electrospray Ionization (ESI) high-resolution mass spectrometry was carried on a Waters XEVO-G2 XS Q-TOF mass spectrometer equipped with an electrospray ion source in positive mode (capillary voltage: 3.0 kV, desolvation gas flow: 900 L $h^{-1}$, temperature: 60 °C) with a resolution R = 22,000 using 200 pg μ$L^{-1}$ Leu-Enk (*m/z* = 556.2771) as a "lock mass". Samples were run using 2 mobile phases: A = 0.1% formic acid in $H_2O$ and B = 0.1% formic acid in $CH_3CN$ on a Waters Acquity UPLC BEH C18 column (2.1 × 50 mm, 1.7 μm); flow rate = 0.6 mL $min^{-1}$, runtime = 3.00 min, column T = 60 °C, mass detection: 50–1500 Da. Gradient: 0–0.15 min: 2% B; 0.15–1.85 min: 2% → 100% B; 1.85–2.05: 100% B; 2.05–2.10 min: 100% → 2% B; 2.10–3.00 min: 100% B. Data processing was performed using Waters MassLynx Mass Spectrometry Software 4.1.

HPLC purifications were performed on a Waters preparative automated HPLC with mass detection. Samples were run using 3 mobile phases: A = $H_2O$, B = $CH_3CN$ and C = 1% 4M $NH_4OH$ in $CH_3CN$ on a Xbridge PREP C18 column (5 μm 19 × 150 mm). Flowrate = 30 mL $min^{-1}$. Gradient: 0 – 2.5 min: 95% A, 5% B; 2.5 – 17.5 min: 5 → 40% B; 17.5 – 20.90 min: 40 → 95% B; 20.90 – 21.00 min: 95 → 5% B; 1 mL $min^{-1}$ C was mixed throughout the whole run. Fractions containing the product were automatically collected based on observed mass (detection range 100–1500 Da) and UV-signal after which they were lyophilized to obtain the pure products.

**Synthetic procedures**

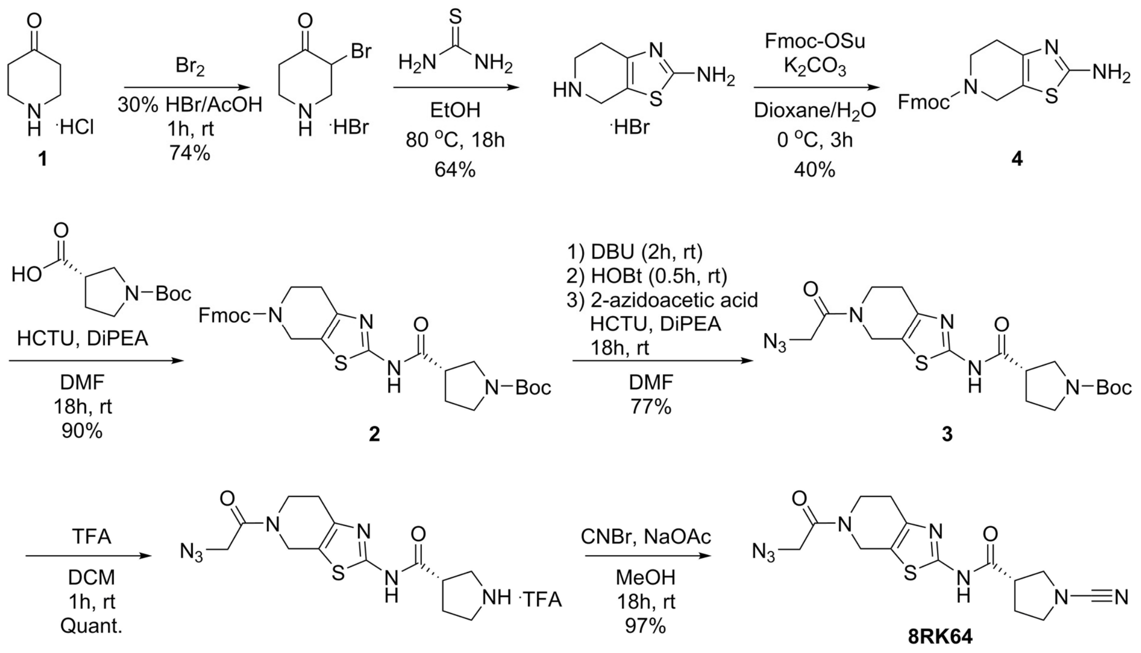


**(9*H*-fluoren-9-yl)methyl 2-amino-6,7-dihydrothiazolo[5,4-*c*]pyridine-5(4*H*)-carboxylate (4).** Step 1: 4-Piperidone monohydrate hydrochloride (10.0 g, 65.1 mmol, 1.0 eq.) was dissolved in AcOH (50 mL). 33% HBr in AcOH (0.82 mL) and bromine (1.67 mL, 32.55 mmol, 0.5 eq.) were added at rt. The resulting reaction mixture was stirred at rt for 1 hour. The reaction mixture was concentrated *in vacuo* and the solids (white/yellow) were suspended in acetone (100 mL). The suspension was refluxed at 63 °C for 2 hours. The solid material was collected in a glass filter and the solids were washed with acetone under reduced pressure to obtain 3-bromopiperidin-4-one-hydrobromide as an off-white powder (12.5 g, 48.3 mmol). This material was not further purified and was used as such.

<u>*Step 2*</u>: 3-bromopiperidin-4-one-hydrobromide (12.5 g, 48.3 mmol, 1.0 eq.) was dissolved in EtOH (150 mL) and thiourea (3.67 g, 48.3 mmol, 1.0 eq.) was added. The resulting reaction mixture was heated at 80 °C for 18 hours. The formed precipitates were collected in a glass filter and were washed with EtOH under reduced pressure to yield 4,5,6,7-tetrahydrothiazolo[5,4-*c*]pyridine-2-amine hydrobromide as an off-white powder (7.4 g, 31.1 mmol).

<u>*Step 3:*</u> To a solution of 4,5,6,7-tetrahydrothiazolo[5,4-*c*]pyridine-2-amine hydrobromide (7.34 g, 31.1 mmol, 1.0 eq.) in 1,4-dioxane (45 mL) were added $H_2O$ (75 mL) and $K_2CO_3$ (8.59 g, 62.1 mmol, 2.0 eq.). The resulting reaction mixture was cooled to 0 °C and Fmoc-OSu (15.72 g, 46.6 mmol, 1.5 eq.) was added. The reaction mixture was allowed to warm to rt and was stirred for 18 hours. The reaction mixture was poured into $H_2O$ (500 mL) and the aqueous layer was extracted 3× with DCM. The combined organic layers were washed with BRINE (500 mL), dried over $Na_2SO_4$, filtered and concentrated. The resulting residue was purified by Büchi flash chromatography (DCM → 4% MeOH/DCM) to yield the titled compound as a white solid (4.43 g, 11.7 mmol, 38%). $^1$H NMR (300 MHz, DMSO-$d_6$) δ 7.90 (d, $J$ = 7.4 Hz, 2*H*), 7.63 (s, 2*H*), 7.48 – 7.27 (m, 4*H*), 6.82 (s, 2*H*), 4.48 – 4.25 (m, 5*H*), 3.68 – 3.42 (m, 2*H*), 2.47 – 2.24 (m, 2*H*). HR-MS calculated for $C_{21}H_{19}N_3O_2S$ $[M+H]^+$ 378.1276, found 378.1269.

**(9*H*-fluoren-9-yl)methyl (*S*)-2-(1-(*tert*-butoxycarbonyl)pyrrolidine-3-carboxamido)-6,7-dihydrothiazolo[5,4-*c*]pyridine-5(4*H*)-carboxylate (2).** To a solution of (*S*)-1-Boc-1-pyrrolidine-3-carboxylic acid (3.4 g, 15.8 mmol, 1.5 eq) in DMF (80 mL) was added HCTU (6.54 g, 15.8 mmol, 1.5 eq) and DiPEA (5.5 mL, 31.6 mmol, 3.0 eq.) and the resulting reaction mixture was stirred for 10 min. at rt. Compound **4** (3.98 mmol, 10.5 mmol, 1.0 eq.) was added and the reaction mixture was stirred at rt for 18 hours. The solvents were evaporated under reduced pressure. The crude material was taken up in EtOAc (200 mL) and the organic layer was washed with 1M HCl (2× 100 mL), sat. aq. $NaHCO_3$ (2× 100 mL) and BRINE (100 mL). The organic layer was dried over $Na_2SO_4$, filtered and concentrated. The resulting residue was purified by Büchi flash chromatography (DCM → 4% MeOH/DCM) to yield **2** as a white foam

(5.44 g, 9.5 mmol, 90%). $^1$H NMR (300 MHz, $CDCl_3$-*d*) δ 9.82 (s, 1*H*), 7.77 (s, 2*H*), 7.58 (s, 2*H*), 7.41 (t, *J* = 7.4 Hz, 2*H*), 7.32 (t, *J* = 7.4 Hz, 2H), 4.72 – 4.45 (m, 4*H*), 4.29 (t, *J* = 6.6 Hz, 1*H*), 3.85 – 3.51 (m, 5*H*), 3.51 – 3.35 (m, 1*H*), 3.13 (s, 1*H*), 2.69 (s, 2*H*), 2.36 – 2.12 (s, 2*H*), 1.49 (s, 9*H*). $^{13}$C NMR (75 MHz, $CDCl_3$-*d*) δ 170.2, 156.2, 155.3, 154.3, 143.8, 141.3, 127.8, 127.1, 124.9, 120.0, 79.8, 67.5, 48.3, 47.3, 45.4, 41.7, 28.5. HR-MS calculated for $C_{31}H_{34}N_4O_5S$ $[M+H]^+$ 575.2328, found 575.2336.

***Tert*-butyl(*S*)-3-((5-(2-azidoacetyl)-4,5,6,7-tetrahydrothiazolo[5,4-*c*]pyridin-2-yl) carbamoyl) pyrrolidine-1-carboxylate (3).** To a solution of **2** (500 mg, 0.87 mmol, 1.0 eq.) in DMF (10 mL) was added 1,8-Diazabicyclo[5.4.0]undec-7-ene (65 µL, 0.44 mmol, 0.5 eq.). The resulting reaction mixture was stirred at rt for 2.5 hours. Complete Fmoc removal was confirmed by TLC and LC-MS. 1-Hydroxybenzotriazole hydrate (176 mg, 1.3 mmol, 1.5 eq.) was added and the reaction mixture was stirred at rt for 30 min. In a separate round bottom flask 2-Azidoacetic acid (228 µL, 3.0 mmol, 3.5 eq.) was dissolved in DMF (5 mL) and HCTU (1.26 g, 3.0 mmol, 3.5 eq.) and DiPEA (758 µL, 4.4 mmol, 5.0 eq.) were added. The resulting reaction mixture was stirred at rt for 10 min. This reaction mixture was added to the first reaction mixture and the whole was stirred at rt for 18 hours. The reaction mixture was concentrated *in vacuo* and the crude material was taken up in EtOAc (100 mL). The organic layer was washed with 1M HCl (2× 100 mL), sat. aq. $NaHCO_3$ (2× 100 mL) and BRINE (100 mL). The 1M HCl phase was extracted with EtOAc (50 mL) and the sat. aq. $NaHCO_3$ phase was extracted with EtOAc (50 mL). All organic layers were combined, dried over $Na_2SO_4$, filtered and concentrated. The resulting residue was purified by Büchi flash chromatography (DCM → 5% MeOH/DCM) to yield **3** as a white foam (292 mg, 0.67 mmol, 77%). $^1$H NMR (300 MHz, $CDCl_3$-*d*) (Mixture of rotamers) δ 4.70 (s, 1*H*), 4.47 (s, 1*H*), 3.99 (d, *J* = 8.8 Hz, 2*H*), 3.92 – 3.82 (m, 1*H*), 3.74 – 3.42 (m, 4*H*), 3.32 (dt, *J* = 10.7, 7.7 Hz, 1*H*), 3.20 – 3.03 (m, 1*H*), 2.78 – 2.70 (m, 1*H*), 2.70 – 2.61 (m, 1*H*), 2.29 – 2.04 (m, 2*H*), 1.38 (s, 9*H*). $^{13}$C NMR (75 MHz, $CDCl_3$-*d*) (Mixture of rotamers) δ 166.4, 166.3, 156.6, 154.3, 143.8, 141.6, 119.4, 117.6, 79.8, 51.1, 51.0, 48.4, 45.4, 44.5, 43.6, 42.9, 42.7, 40.4, 40.2, 28.9, 28.5, 27.1, 26.2. HR-MS calculated for $C_{18}H_{25}N_7O_4S$ $[M+H]^+$ 436.1767, found 436.1755.

**(*S*)-*N*-(5-(2-azidoacetyl)-4,5,6,7-tetrahydrothiazolo[5,4-*c*]pyridin-2-yl)-1-cyanopyrrolidine-3-carboxamide (8RK64).** *Step 1:* To a solution of **3** (40.0 mg, 92 µmol, 1.0 eq.) in DCM (1.0 mL) was added TFA (1.0 mL). The resulting reaction mixture was stirred at rt for 2 hours. The solvents were evaporated under reduced pressure and the residue was co-evaporated with DCM (3× 5.0 mL). This material was not further purified and was used as such. _Step 2:_ The TFA-salt from Step 1 was dissolved in MeOH (1.0 mL) and NaOAc (38 mg, 0.46 mmol, 5.0 eq.) and cyanogen bromide (39 mg, 0.37 mmol, 4.0 eq.) were added. The resulting reaction mixture was stirred at rt for 18 hours. The solvents were evaporated under reduced pressure and the crude material was taken up in EtOAc (25 mL). The organic layer was washed with sat. $NaHCO_3$ (2× 25 mL) and BRINE (25 mL). The organic layer was dried over $Na_2SO_4$, filtered and concentrated. The resulting residue was purified by Büchi flash chromatography (DCM → 5% MeOH/DCM) to yield **8RK64** as a white foam (32 mg, 89 µmol, 97%). $^1$H NMR (300 MHz, $CDCl_3$-*d*) (Mixture of rotamers) δ 4.80 (s, 1*H*), 4.58 (s, 1*H*), 4.10 (d, *J* = 10.4 Hz, 2*H*), 4.01 – 3.94 (m, 1*H*), 3.75 (s, 1*H*), 3.72 (s, 2*H*), 3.69 – 3.60 (m, 1*H*), 3.59 – 3.47 (m, 1*H*), 3.39 – 3.23 (m, 1*H*), 2.89 – 2.81 (m, 1*H*), 2.81 – 2.72 (m, 1*H*), 2.31 (q, *J* = 7.2 Hz, 2*H*). $^{13}$C NMR (75 MHz, $CDCl_3$-*d*) (Mixture of rotamers) δ 169.5, 169.2, 166.5, 166.4, 156.4, 156.3, 143.8, 141.7, 119.6, 117.9, 116.8, 52.6, 51.2, 51.0, 50.3, 44.2, 42.9, 42.7, 40.4, 40.3, 29.5. HR-MS calculated for $C_{14}H_{16}N_8O_2S$ $[M+H]^+$ 361.1195, found 361.1197.

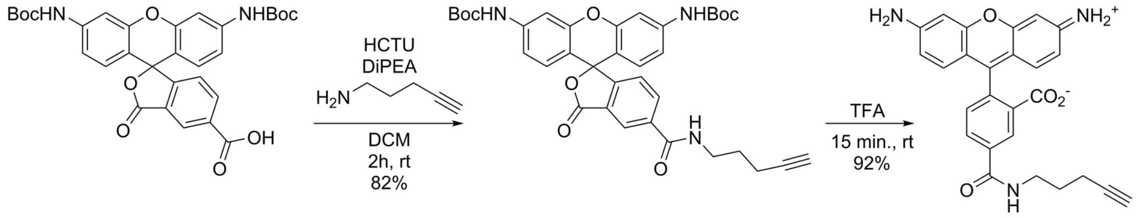


5

6

**Di-*tert*-butyl (3-oxo-5-(pent-4-yn-1-ylcarbamoyl)-3*H*-spiro[isobenzofuran-1,9'-xanthene]-3',6'-diyl)dicarbamate (5).** 3',6'-bis((*tert*-butoxycarbonyl)amino)-3-oxo-3*H*-spiro[isobenzofuran-1,9'-xanthene]-5-carboxylic acid[1] (127 mg, 0.22 mmol, 1.0 eq.) was dissolved in DCM (5 mL). To this were added DiPEA (153 µL, 0.88 mmol, 4.0 eq.), HCTU (137 mg, 0.33 mmol, 1.5 eq.) and pent-4-yn-1-amine (34 µL, 0.33 mmol, 1.5 eq.) and the mixture was stirred at rt until TLC indicated a complete conversion of starting material after 2 hours. The mixture was diluted with DCM (50 mL) and extracted with 1M HCl (2× 50 mL), sat. aq. $NaHCO_3$ (2× 50 mL) and BRINE (50 mL), dried over $Na_2SO_4$, and concentrated under reduced pressure. The crude material was purified by Büchi flash chromatography (5 → 50% EtOAc/*n*-hept.) and the product was obtained as a colorless solid (115 mg, 0.18 mmol, 82%).

$^1$H NMR (300 MHz, $CDCl_3$-*d*) δ 8.4 (d, 1*H*), 8.2 (dd, *J* = 8.0, 1.6 Hz, 1*H*), 7.5 (d, *J* = 2.2 Hz, 2*H*), 7.2 (t, *J* = 5.8 Hz, 1*H*), 7.1 (dd, *J* = 8.0, 0.7 Hz, 1*H*), 7.0 (s, 2*H*), 6.9 (dd, *J* = 8.7, 2.2 Hz, 2*H*), 6.6 (d, *J* = 8.6 Hz, 2*H*), 3.6 (q, *J* = 6.5 Hz, 2*H*), 2.34 – 2.24 (m, 2*H*), 2.0 (t, *J* = 2.6 Hz, 1*H*), 1.9 (q, *J* = 6.8 Hz, 2*H*), 1.5 (s, 18*H*). $^{13}$C NMR (75 MHz, $CDCl_3$-*d*) δ 168.9, 165.8, 155.4, 152.4, 151.8, 140.9, 136.6, 134.7, 128.2, 126.7, 124.3, 123.2, 114.3, 112.1, 106.2, 83.5, 81.1, 69.4, 39.6, 28.2, 27.8, 16.2. HR-MS calculated for $C_{36}H_{37}N_3O_8$ [M+H]$^+$ 640.2659, found 640.2654.

**2-(6-amino-3-iminio-3*H*-xanthen-9-yl)-5-(pent-4-yn-1-ylcarbamoyl)benzoate (6).** Compound **5** (115 mg, 0.18 mmol) was dissolved in DCM (3 mL) after which TFA (3 mL) was added. The mixture was stirred at rt until TLC and LC-MS analysis indicated a complete reaction. Toluene (15 mL) was added and the reaction was concentrated under reduced pressure, followed by co-evaporation with toluene (2×). The product was dissolved in $H_2O/CH_3CN$/AcOH 1:1:0.1 v/v/v (10 mL) and lyophilized, which yielded the product as a red/brown solid (91 mg, 0.16 mmol, 92%). $^1$H NMR (300 MHz, MeOD-$d_4$) δ 8.76 (d, *J* = 1.8 Hz, 1*H*), 8.24 (dd, *J* = 7.9, 1.9 Hz, 1*H*), 7.50 (d, *J* = 8.0 Hz, 1*H*), 7.03 (d, *J* = 9.6 Hz, 2*H*), 6.79 (m, 4*H*), 3.66 – 3.49 (m, 2*H*), 2.42 – 2.21 (m, 3*H*), 1.98 – 1.92 (m, 2*H*). $^{13}$C NMR (75 MHz, MeOD-$d_4$) δ 168.3, 167.5, 161.2, 161.0, 159.6, 138.1, 137.6, 132.9, 132.7, 132.2, 131.9, 131.2, 117.9, 114.7, 98.4, 84.2, 70.1, 40.4, 29.4, 16.9. HR-MS calculated for $C_{26}H_{21}N_3O_4$ [M+H]$^+$ 440.1610, found 440.1627.

**General procedure for click chemistry**

Compound **8RK64** (1.0 eq.) and the appropriate alkyne (1.2 eq.) were dissolved in dry DMF (1-2 mL). Argon was bubbled through the reaction mixture for 30 min. An aqueous solution of $CuSO_4 \cdot 5H_2O$ (100 µL, 0.50 eq.) and an aqueous solution of sodium ascorbate (100 µL, 0.75 eq.) were added. The aqueous solutions of sodium ascorbate and $CuSO_4.5H_2O$ were prepared in 5.0 mL volume and degassed for 30 min. with argon bubbling. After addition of sodium ascorbate and $CuSO_4 \cdot 5H_2O$ the resulting reaction mixture was stirred at rt for 2-18 hours, concentrated under reduced pressure and purified as described. In case necessary, excess alkyne was reacted with 2-Azidoacetic acid for ease of purification.

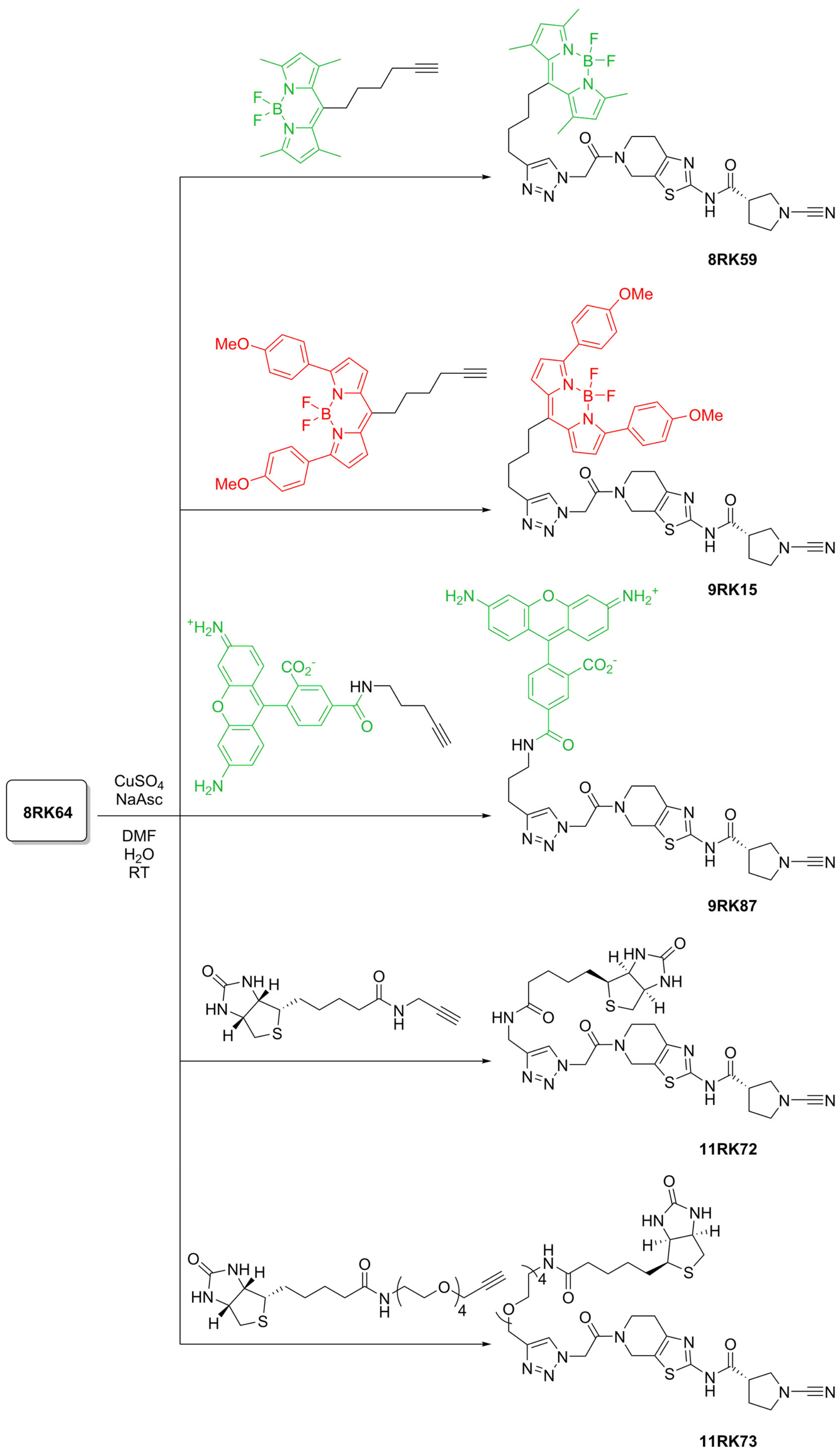

8RK64
CuSO4
NaAsc
DMF
H2O
RT
8RK59
9RK15
9RK87
11RK72
11RK73

**BodipyFL probe 8RK59.** Synthesized according to the general procedure for click chemistry using **8RK64** (25.0 mg, 69 µmol) and BodipyFL-alkyne[2] (27.3 mg, 83 µmol). The crude material was taken up in DCM (20 mL). The organic layer was washed with BRINE (10 mL), dried over $Na_2SO_4$, filtered and concentrated. The resulting residue was purified by Büchi flash chromatography (DCM → 5% MeOH/DCM) to yield **8RK59** as a red solid (10 mg, 14 µmol, 21%). $^1$H NMR (300 MHz, DMSO-$d_6$) (Mixture of rotamers) δ 12.37 – 12.22 (m, 1*H*), 7.81 (d, *J* = 1.9 Hz, 1*H*), 6.26 (s, 2*H*), 5.54 (d, *J* = 20.1 Hz, 2*H*), 4.73 (d, *J* = 36.4 Hz, 2*H*), 3.85 (m, 2*H*), 3.72 – 3.43 (m, 4*H*), 3.07 – 2.95 (m, 2*H*), 2.89 – 2.67 (m, 4*H*), 2.43 (m, 12*H*), 2.30 – 2.01 (m, 3*H*), 1.87 (m, 2*H*), 1.67 (d, 2*H*). $^{13}$C NMR (75 MHz, DMSO-$d_6$) (Mixture of rotamers) δ 170.8, 165.5, 165.3, 157.6, 156.5, 153.5, 147.1, 146.6, 143.6, 143.2, 141.3, 131.1, 124.2, 122.1, 118.6, 118.3, 117.5, 52.5, 51.3, 51.1, 50.4, 48.5, 43.5, 42.6, 42.2, 31.2, 29.9, 29.7, 28.1, 27.1, 26.3, 25.0, 16.3, 14.4. HR-MS calculated for $C_{33}H_{39}BF_2N_{10}O_2S$ $[M-F]^+$ 669.3061, found 669.3063.

**BodipyTMR probe 9RK15.** Synthesized according to the general procedure for click chemistry using **8RK64** (24.0 mg, 67 µmol) and BodipyTMR-alkyne[2] (39.0 mg, 81 µmol). The crude material was taken up in EtOAc (30 mL). The organic layer was washed with $H_2O$ (10 mL), dried over $Na_2SO_4$, filtered and concentrated. The resulting residue was purified by Büchi flash chromatography (DCM → 5% MeOH/DCM) to yield **9RK15** as a red solid (17.7 mg, 21 µmol, 31%). $^1$H NMR (300 MHz, DMSO-$d_6$) (Mixture of rotamers) δ 12.27 (d, *J* = 10.7 Hz, 1*H*), 7.76 – 7.86 (m, 5*H*), 7.68 – 7.62 (m, 2*H*), 7.05 – 6.98 (m, 4*H*), 6.80 (d, *J* = 4.3 Hz, 2*H*), 5.51 (d, *J* = 20.0 Hz, 2*H*), 4.70 (d, *J* = 34.1 Hz, 2*H*), 3.85 – 3.77 (m, 8*H*), 3.63 – 3.57 (m, 1*H*), 3.57 – 3.49 (m, 1*H*), 3.49 – 3.40 (m, 2*H*), 3.16 – 3.03 (m, 2*H*), 2.85 – 2.77 (s, 1*H*), 2.77 – 2.61 (m, 3*H*), 2.30 – 1.97 (m, 3*H*), 1.79 (m, 2.30 – 1.97, 4*H*). $^{13}$C NMR (75 MHz, DMSO-$d_6$) (Mixture of rotamers) δ 170.9, 170.8, 165.5, 165.3, 160.7, 157.0, 156.6, 156.5, 146.7, 146.6, 143.6, 143.2, 136.2, 131.2, 125.1, 124.1, 120.7, 118.6, 118.3, 117.5, 114.2, 73.5, 72.7, 70.2, 55.7, 52.5, 51.3, 51.1, 50.4, 43.5, 42.6, 42.2, 33.7, 30.0, 29.7, 29.5, 29.2, 27.1, 26.3, 25.1. HR-MS calculated for $C_{43}H_{43}BF_2N_{10}O_4S$ $[M-F]^+$ 825.3274, found 825.3346.

**Rhodamine probe 9RK87.** Synthesized according to the general procedure for click chemistry using **8RK64** (18.57 mg, 52 µmol) and **6** (39.17 mg, 62 µmol). The crude material was purified by preparative HPLC to yield **9RK87** as a red solid (3.56 mg, 4.45 µmol, 9%). $^1$H NMR (300 MHz, DMSO-$d_6$) (Mixture of rotamers) δ 12.27 (m, 1*H*), 8.88 (t, *J* = 5.6 Hz, 1*H*), 8.44 (s, 1*H*),

8.24 (dd, *J* = 8.0, 1.5 Hz, 1*H*), 7.82 (d, *J* = 2.9 Hz, 1*H*), 7.33 (d, *J* = 8.0 Hz, 1*H*), 6.44 – 6.26 (m, 5*H*), 5.73 – 5.47 (m, 5*H*), 4.71 (d, *J* = 37.0 Hz, 2*H*), 3.83 (d, *J* = 5.3 Hz, 2*H*), 3.68 – 3.57 (m, 1*H*), 3.54 (dd, *J* = 6.1, 3.7 Hz, 1*H*), 3.48 – 3.36 (m, 5*H*), 2.86 – 2.78 (m, 1*H*), 2.78 – 2.62 (m, 3*H*), 2.25 – 2.12 (m, 1*H*), 2.12 – 1.99 (m, 1*H*), 1.98 – 1.84 (m, 2*H*). HR-MS calculated for $C_{40}H_{37}N_{11}O_6S$ $[M+H]^+$ 800.2727, found 800.2749.

**Biotin probe 11RK72.** Synthesized according to the general procedure for click chemistry using **8RK64** (13.0 mg, 36 µmol) and Biotin-alkyne[3] (12.20 mg, 43 µmol). The crude material was purified by preparative HPLC to yield **11RK72** as a white solid (7.23 mg, 11.3 µmol, 31%). HR-MS calculated for $C_{27}H_{35}N_{11}O_4S_2$ $[M+H]^+$ 642.2393, found 642.2393.

**Biotin-PEG$_4$ probe 11RK73.** Synthesized according to the general procedure for click chemistry using **8RK64** (13.0 mg, 36 µmol) and BiotinPEG$_4$-alkyne (19.8 mg, 43 µmol). The crude material was purified by preparative HPLC to yield **11RK73** as a white solid (5.24 mg, 6.4 µmol, 18%). HR-MS calculated for $C_{35}H_{51}N_{11}O_8S_2$ $[M+H]^+$ 818.3442, found 818.3470.

**Synthesis of Ub-Rho-morpholine**

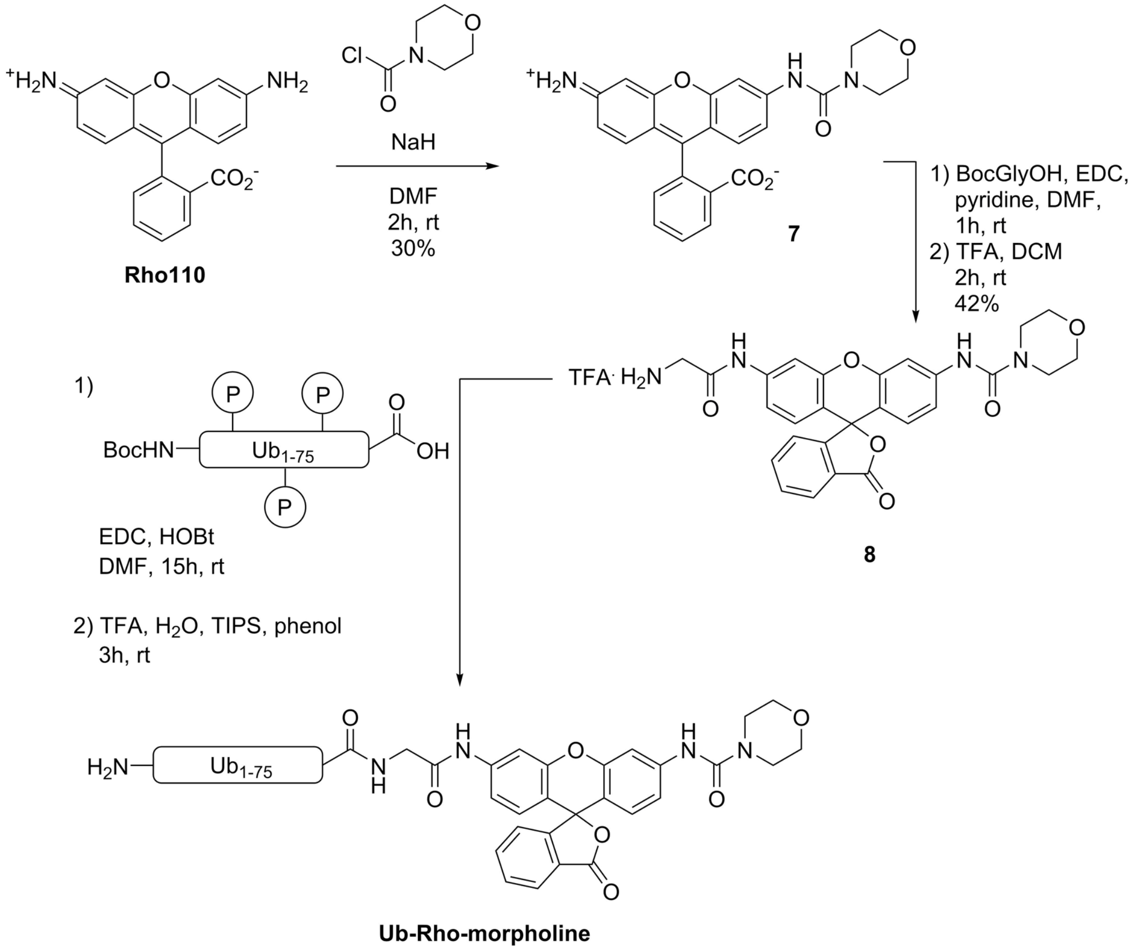


**2-(3-iminio-6-(morpholine-4-carboxamido)-3*H*-xanthen-9-yl)benzoate (7).**[4] Rhodamine110 (500 mg, 1.36 mmol, 1.0 eq.) was dissolved in anhydrous DMF (50 mL) and flushed with argon. NaH (60% w/w in mineral oil, 114 mg, 2.86 mmol, 2.1 eq.) was carefully added in 3 portions and the solution was stirred for 1 hour at rt. 4-Morpholinecarbonyl chloride (156 µL, 1.36 mmol, 1.0 eq.) was added dropwise and the mixture was stirred for another 3 hours. LC-MS analysis indicated the formation of a mixture of unreacted, 1× and 2× morpholinecarbonyl-coupled rhodamine. The mixture was concentrated to dryness under reduced pressure and purified by Büchi flash chromatography (1% AcOH/DCM → 1% AcOH/ (10% MeOH/DCM)) to yield the titled compound as an orange solid (180 mg, 0.40 mmol, 30%). $^1$H NMR (300 MHz, $CDCl_3$) δ 7.94 (d, *J* = 6.5 Hz, 1H), 7.65 – 7.52 (m, 2H), 7.48 (s, 1H), 7.40 (d, *J* = 2.0 Hz, 1H), 7.08 (d, *J* = 6.7 Hz, 1H), 6.85 (dd, *J* = 8.6, 2.1 Hz, 1H), 6.49 (dd, *J* = 12.4, 8.6 Hz, 2H), 6.38 (d, *J* = 2.2 Hz, 1H), 6.27 (dd, *J* = 8.5, 2.2 Hz, 1H), 3.58 (t, *J* = 4.8

Hz, 4H), 3.41 (d, $J$ = 4.9 Hz, 4H). $^{13}$C NMR (75 MHz, $CDCl_3$) δ 170.34, 155.11, 152.90, 151.98, 149.67, 141.76, 135.11, 129.76, 129.04, 128.13, 127.17, 125.03, 124.44, 115.75, 113.04, 111.88, 108.16, 107.67, 101.33, 66.52, 44.32, 43.59.

***N*-(3'-(2-aminoacetamido)-3-oxo-3*H*-spiro[isobenzofuran-1,9'-xanthen]-6'-yl)morpholine-4-carboxamide TFA salt (8).** Compound **7** (158 mg, 0.35 mmol, 1.0 eq.) was dissolved in a mixture of anhydrous DMF (3 mL) and pyridine (2 mL) and flushed with argon. EDC·HCl (135 mg, 0.70 mmol, 2.0 eq.) and 1-Hydroxybenzotriazole hydrate (123 mg, 0.70 mmol, 2.0 eq.) were added and the mixture was stirred at rt for 1 hour after which LC-MS analysis indicated complete conversion. The mixture was concentrated under reduced pressure, dissolved in DCM (20 mL) and extracted with 1M HCl (2× 20 mL) and BRINE (20 mL). The organic layer was dried over $Na_2SO_4$, filtered and concentrated and the resulting white solid was dissolved in a 2:1 v/v mixture of DCM/TFA (5 mL) and stirred for 2 hours at rt until LC-MS analysis indicated complete consumption of starting material. The mixture was concentrated under reduced pressure, co-evaporated with (DCE 3× 20 mL) and purified by Büchi flash chromatography (DCM → 15% MeOH/DCM)) to yield the titled compound as a white solid (74 mg, 0.15 mmol, 42%). $^{1}$H NMR (300 MHz, MeOD-$d_4$) δ 7.99 (dd, $J$ = 7.2, 0.9 Hz, 1H), 7.74 (d, $J$ = 2.0 Hz, 1H), 7.69 (ddd, $J$ = 9.0, 7.2, 1.3 Hz, 2H), 7.48 (d, $J$ = 2.1 Hz, 1H), 7.13 (ddd, $J$ = 8.6, 5.2, 1.5 Hz, 2H), 7.05 (dd, $J$ = 8.7, 2.2 Hz, 1H), 6.67 (d, $J$ = 8.6 Hz, 1H), 6.61 (d, $J$ = 8.7 Hz, 1H), 3.67 (dd, $J$ = 5.7, 3.9 Hz, 4H), 3.49 (dd, $J$ = 5.7, 4.0 Hz, 4H), 3.35 (s, 2H). $^{13}$C NMR (75 MHz, Methanol-$d_4$) δ 171.32, 157.26, 154.32, 152.96, 152.83, 143.63, 141.73, 136.71, 131.23, 129.43, 129.02, 127.61, 125.89, 125.13, 117.24, 116.42, 115.36, 113.85, 108.56, 108.40, 84.47, 67.61, 45.58, 44.85.

**Ub-Rho-morpholine.** Step 1: Sidechain-protected Boc-$Ub_{1-75}$-COOH (~20 µmol) was obtained by Fmoc SPPS on trityl-resin as described in literature.[5] It was dissolved in DMF (4 mL) and to this were added EDC·HCl (10.9 mg, 60 µmol 3.0 eq.), HOBt (7.7 mg, 60 µmol, 3.0 eq.) and compound **8** (28.1 mg, 60 µmol, 3.0 eq.) and the mixture was stirred at rt for 15 hours before being concentrated to dryness under reduced pressure. The resulting residue was dissolved in a mixture of 60:30:10 v/v/v $H_2O/CH_3CN$/AcOH (5 mL) and lyophilized.

Step2: The resulting residue from step 1 was dissolved in a mixture of 90.5/5/2/2.5 v/v/v/v TFA/$H_2O$/TIPS/phenol (5 mL) and stirred for 3 hours at rt. The protein was precipitated from ice-cold $Et_2O$/*n*-pentane (3/1; v/v; 20 mL). The solution was centrifuged and $Et_2O$/*n*-pentane (supernatant) was removed. The pellet was washed with $Et_2O$ (20 mL), the solution was vortexed, the suspension was centrifuged and $Et_2O$ was removed. The wash step was repeated twice. The pellet was dissolved in $H_2O/CH_3CN$/formic acid (65/25/10; v/v/v; 10 mL) and lyophilized. The protein was subsequently purified using RP-HPLC and the product was obtained as a white solid (99.7 mg, 11 μmol, 55%). LC-MS (6 min. run): $R_t$ (min.) 1.87; deconvoluted mass: 8973.0.

#### MS analysis of covalent complex formation between UCHL1 and inhibitors/probes

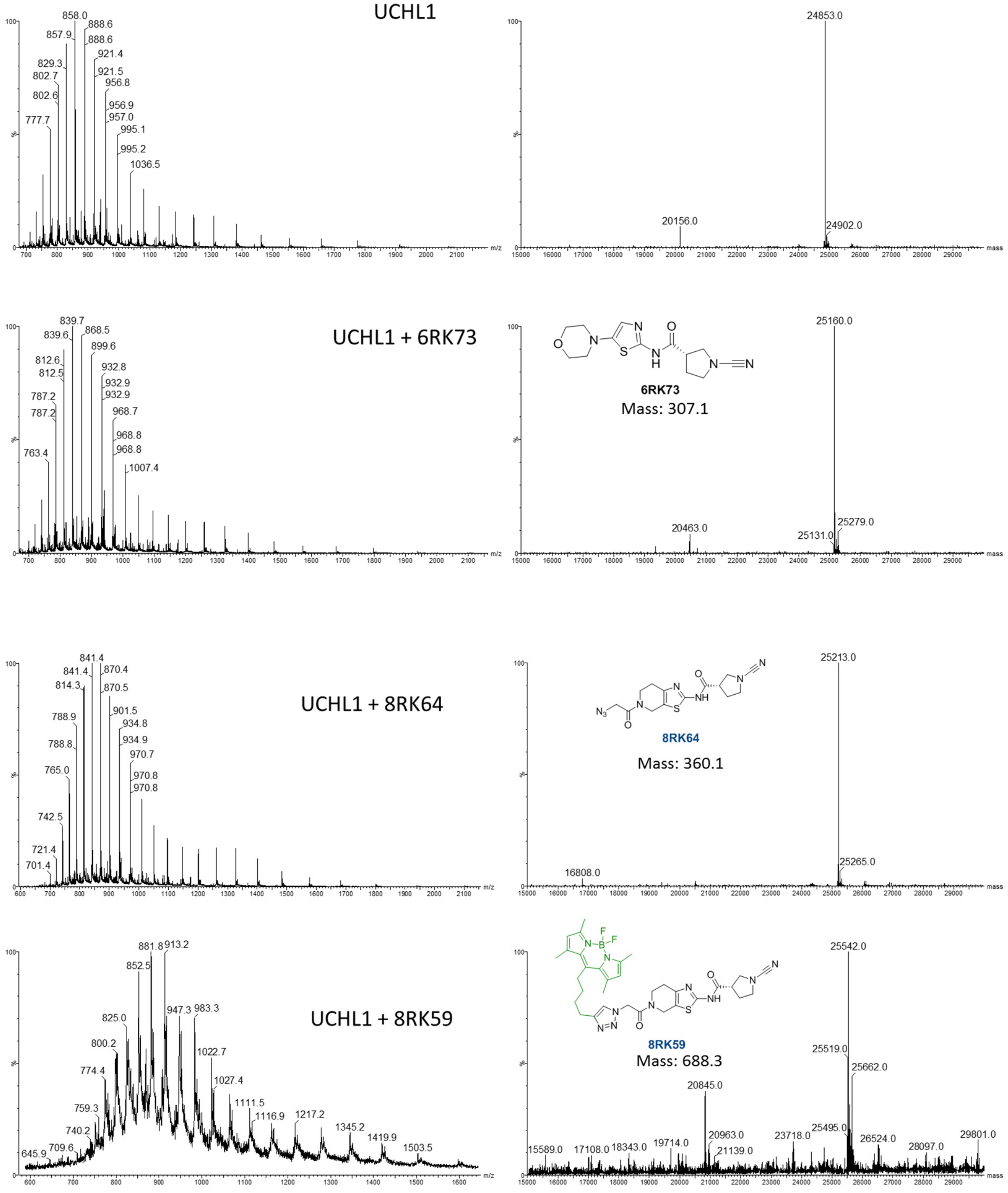

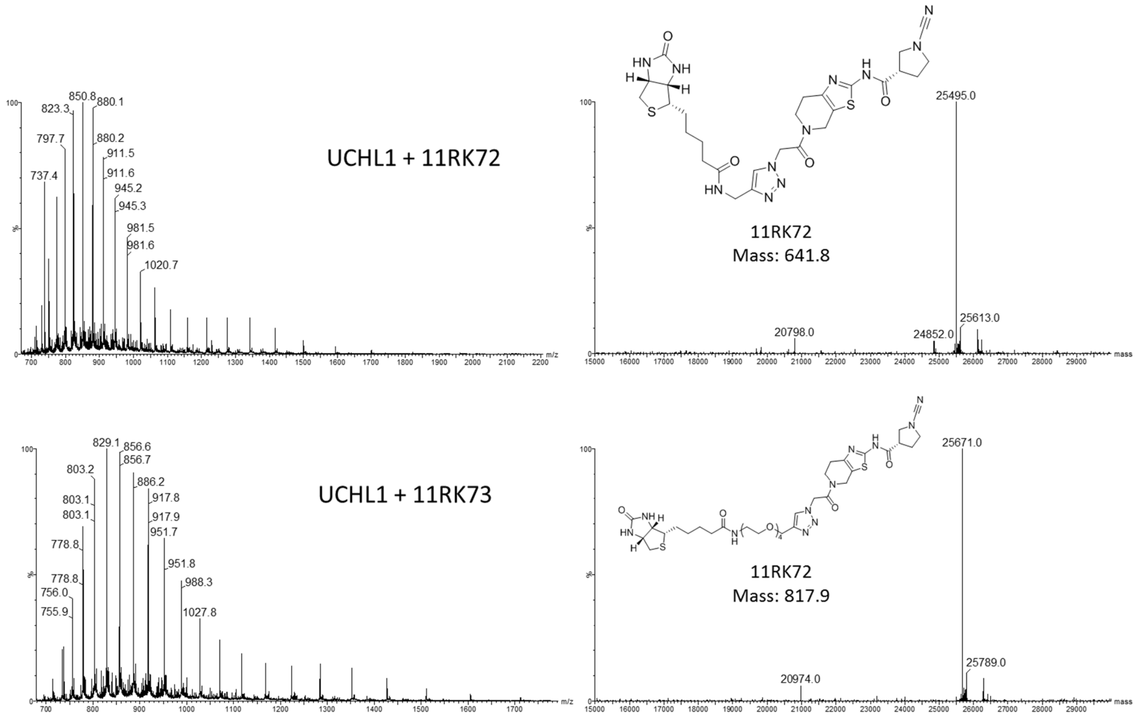
UCHL1 + 11RK72
11RK72
Mass: 641.8
25495.0
20798.0
24852.0
25613.0
UCHL1 + 11RK73
11RK72
Mass: 817.9
25671.0
20974.0
25789.0

#### IC$_{50}$ curves

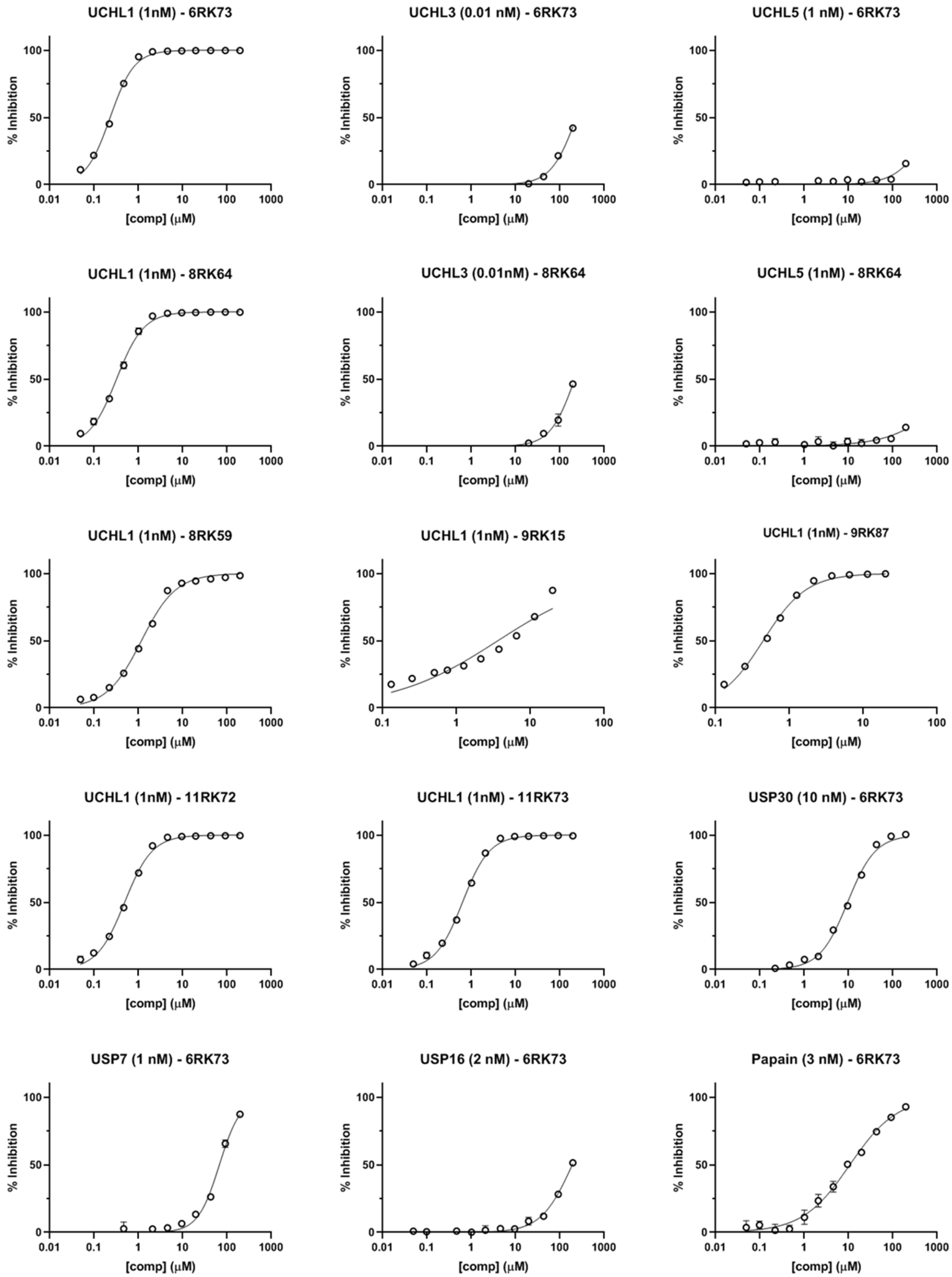
UCHL1 (1nM) - 6RK73
UCHL3 (0.01 nM) - 6RK73
UCHL5 (1 nM) - 6RK73
UCHL1 (1nM) - 8RK64
UCHL3 (0.01nM) - 8RK64
UCHL5 (1nM) - 8RK64
UCHL1 (1nM) - 8RK59
UCHL1 (1nM) - 9RK15
UCHL1 (1nM) - 9RK87
UCHL1 (1nM) - 11RK72
UCHL1 (1nM) - 11RK73
USP30 (10 nM) - 6RK73
USP7 (1 nM) - 6RK73
USP16 (2 nM) - 6RK73
Papain (3 nM) - 6RK73
% Inhibition
[comp] (μM)

### NMR & LC-MS analysis of synthesized compounds

**Compound 4: $^{1}H$ NMR (300.17 MHz, DMSO)**

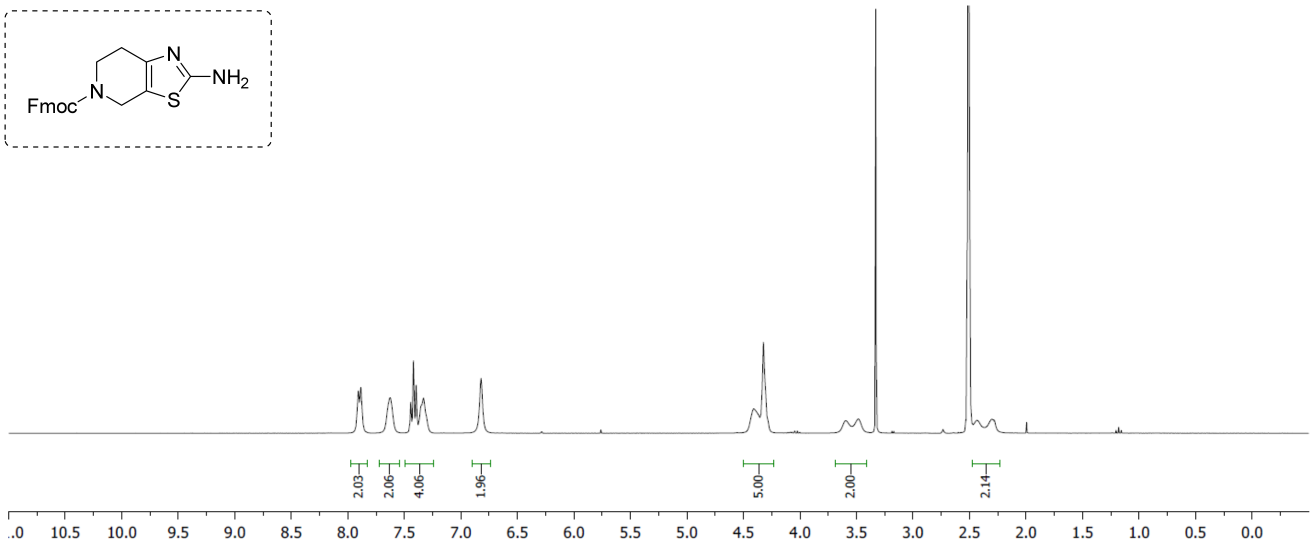


**Compound 4: $^{13}C$ NMR (75.47 MHz, DMSO)**

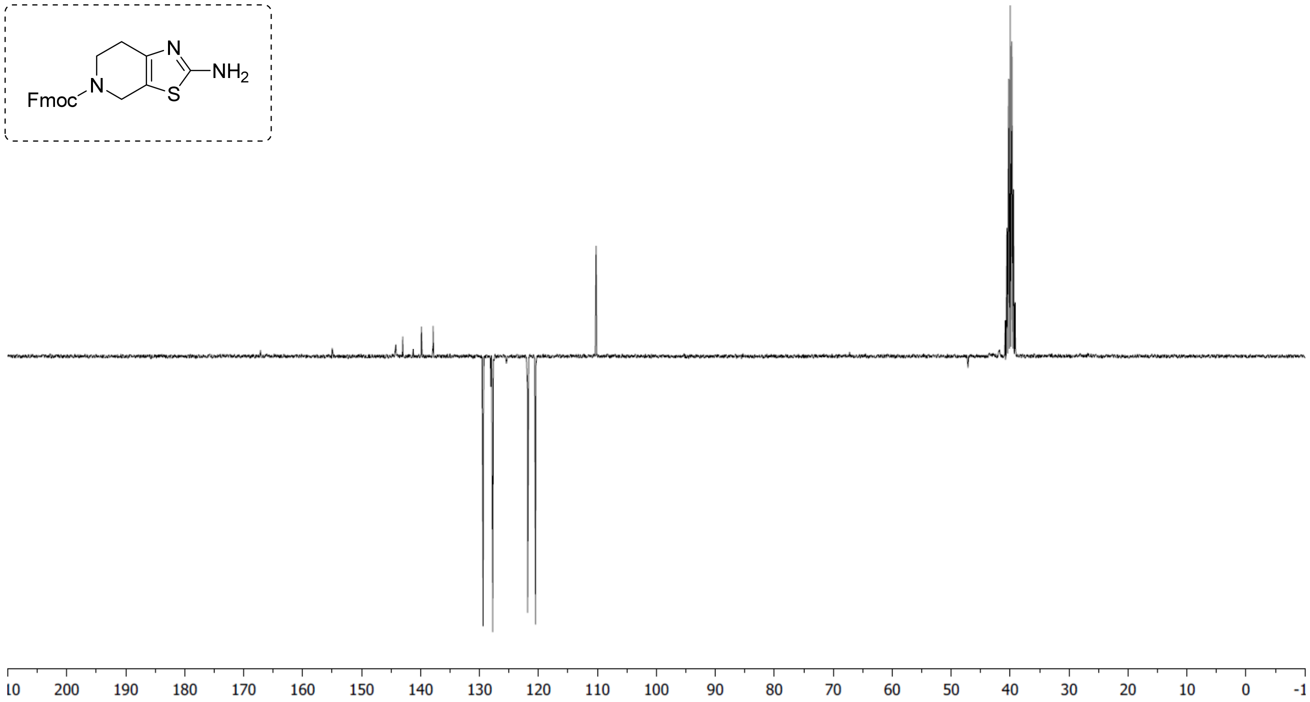

**Compound 2: $^{1}H$ NMR (300.17 MHz, $CDCl_3$)**

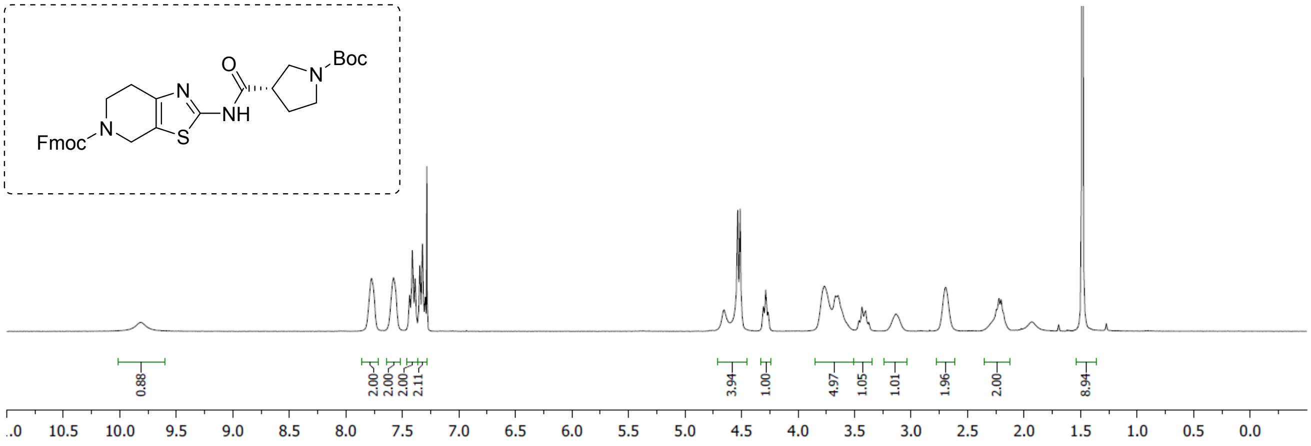


**Compound 2: $^{13}C$ NMR (75.47 MHz, $CDCl_3$)**

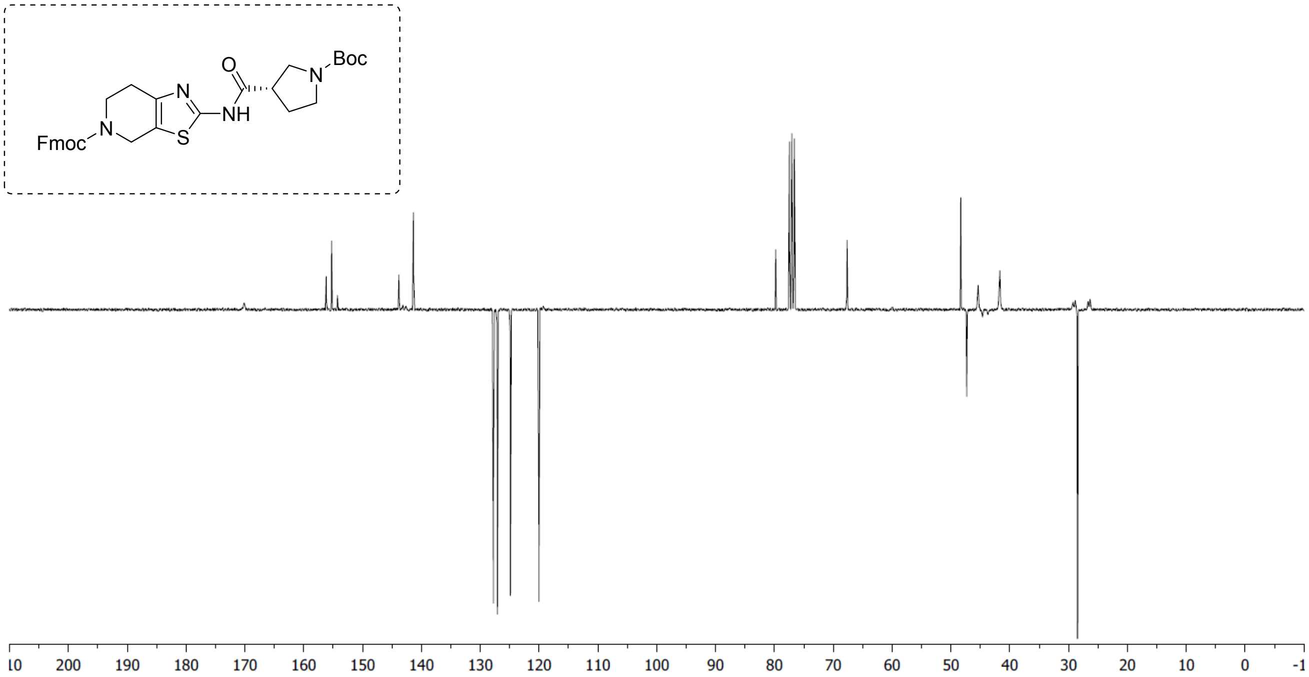


**Compound 3: $^{1}H$ NMR (300.17 MHz, $CDCl_3$)**

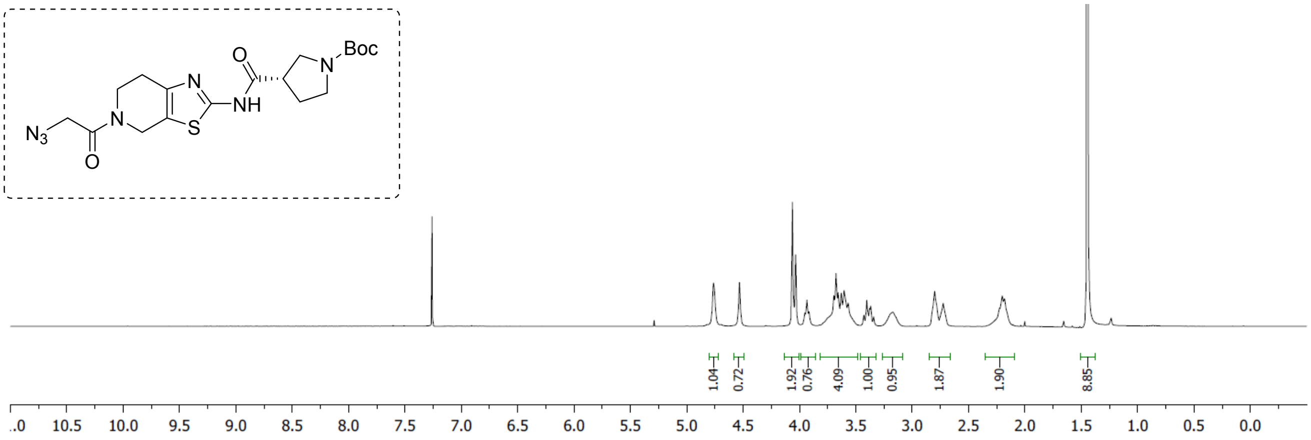

**Compound 3: $^{13}C$ NMR (75.47 MHz, $CDCl_3$)**

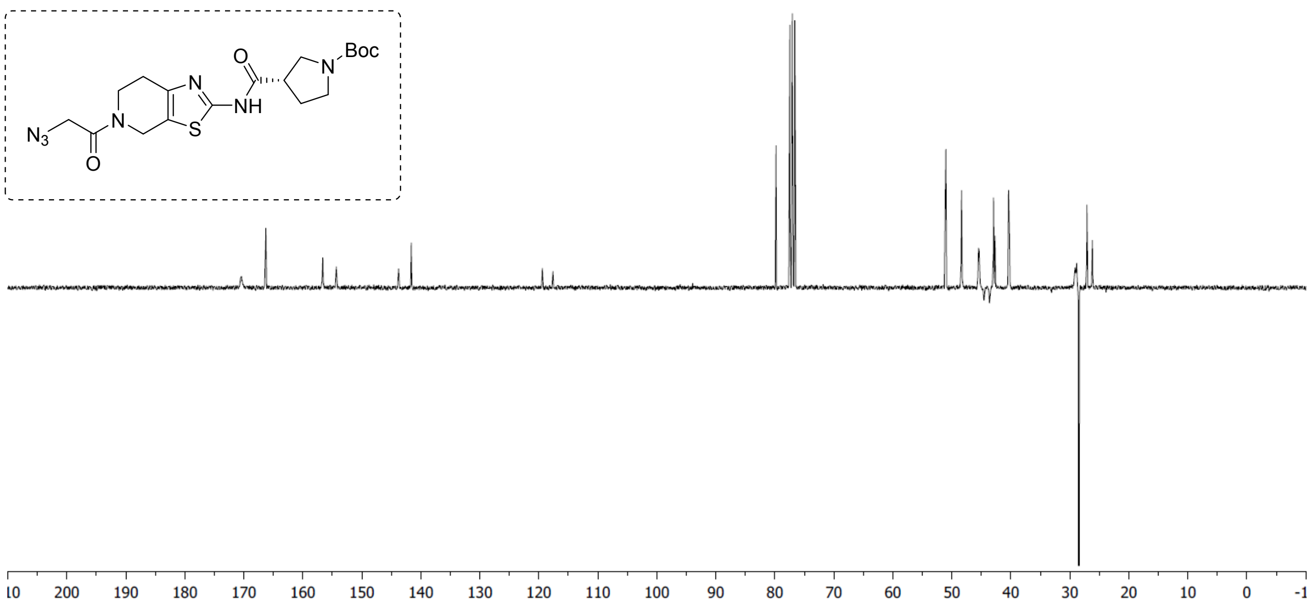


**Compound 8RK64: $^1H$ NMR (300.17 MHz, $CDCl_3$)**

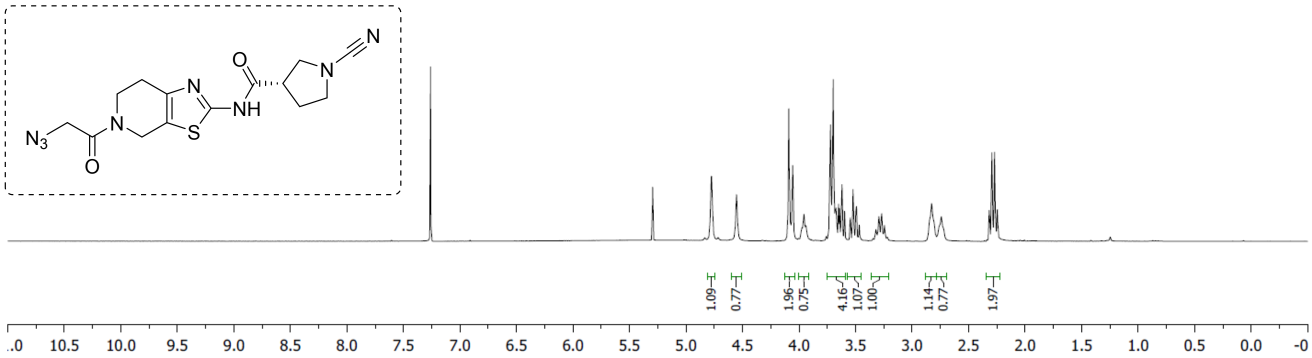


**Compound 8RK64: $^{13}C$ NMR (75.47 MHz, $CDCl_3$)**

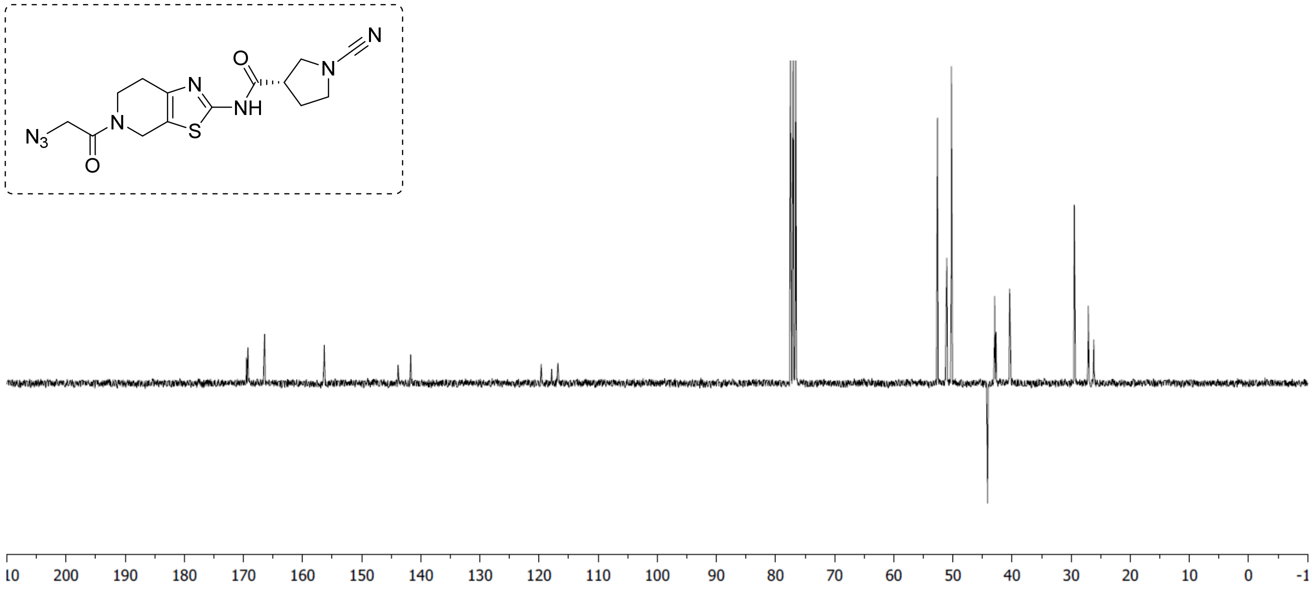

**Compound 8RK59: $^{1}$H NMR (300.17 MHz, DMSO)**

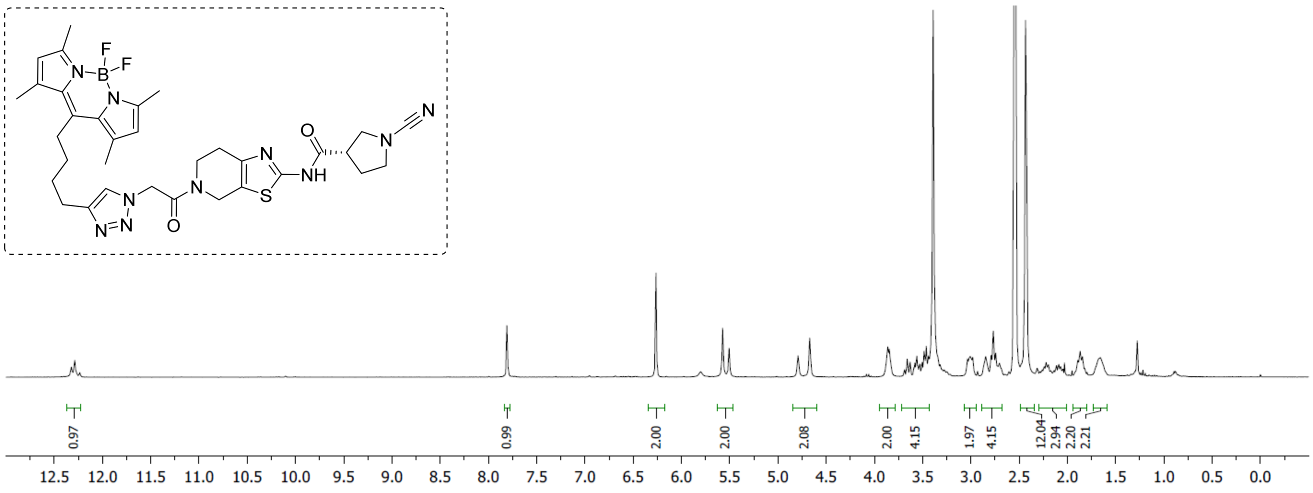


**Compound 8RK59: $^{13}$C NMR (75.47 MHz, DMSO)**

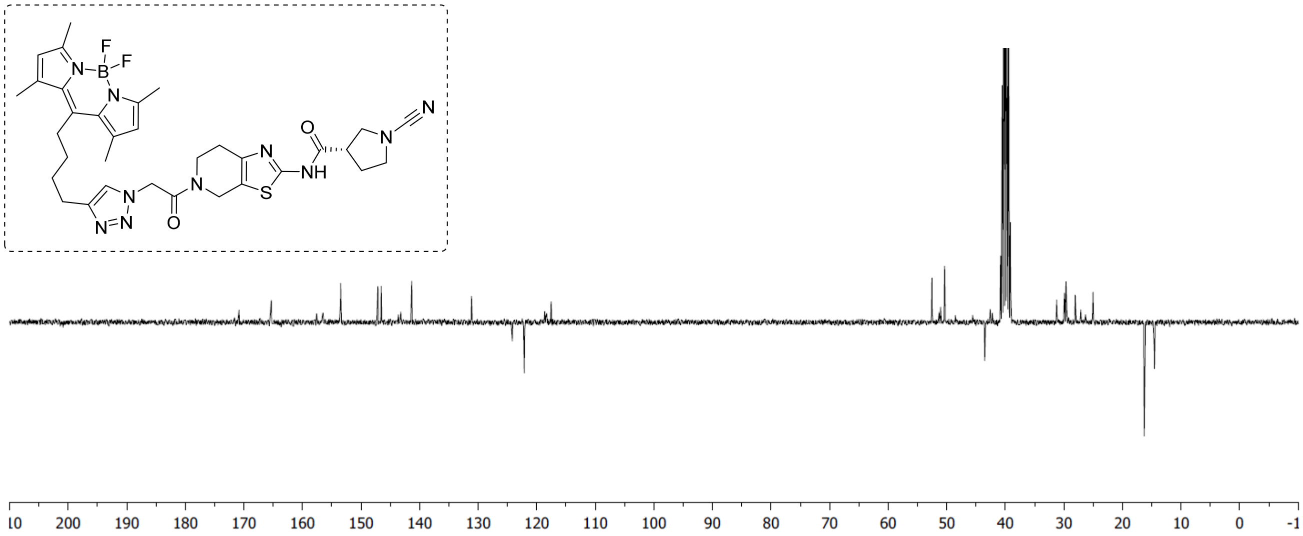


**Compound 9RK15: $^{1}$H NMR (300.17 MHz, DMSO)**

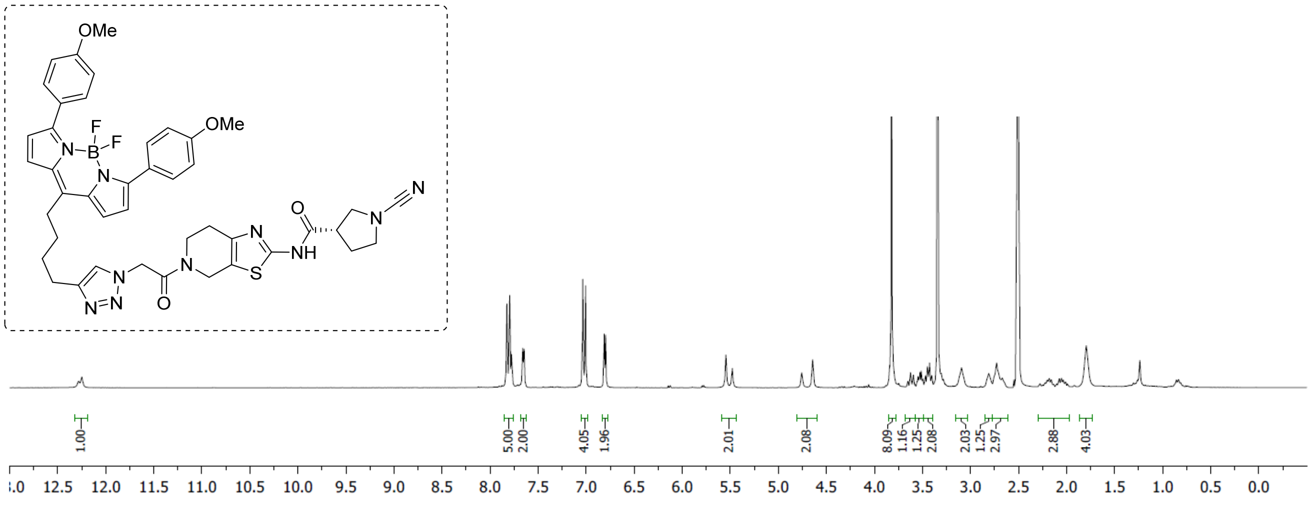

**Compound 9RK15: $^{13}$C NMR (75.47 MHz, DMSO)**

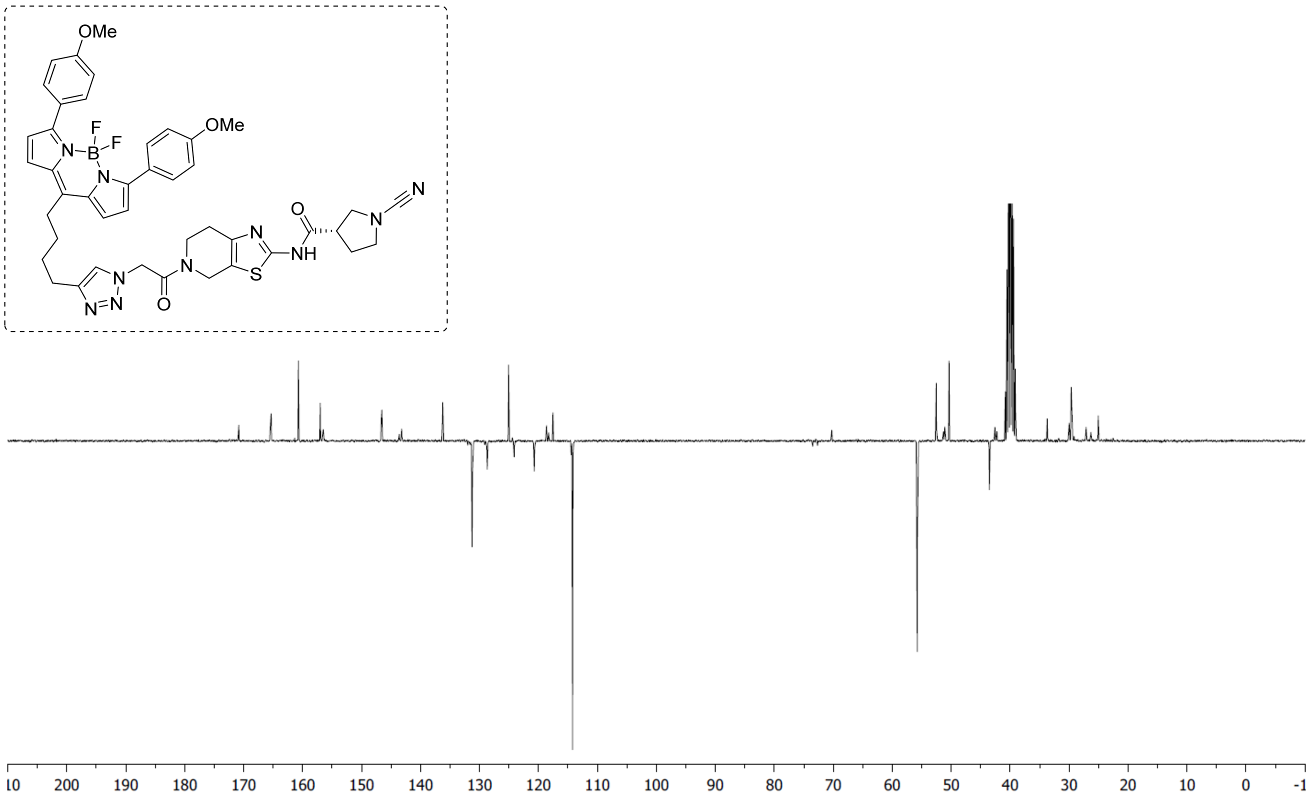


**Compound 5: $^{1}$H NMR (300.17 MHz, CDCl$_3$)**

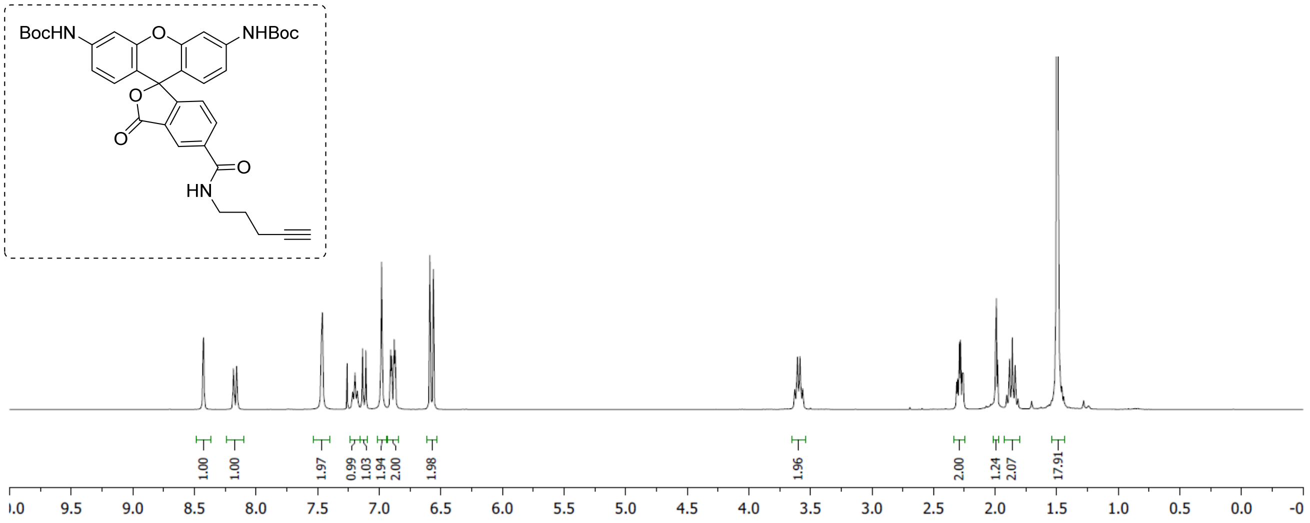

**Compound 5: $^{13}C$ NMR (75.47 MHz, $CDCl_3$)**

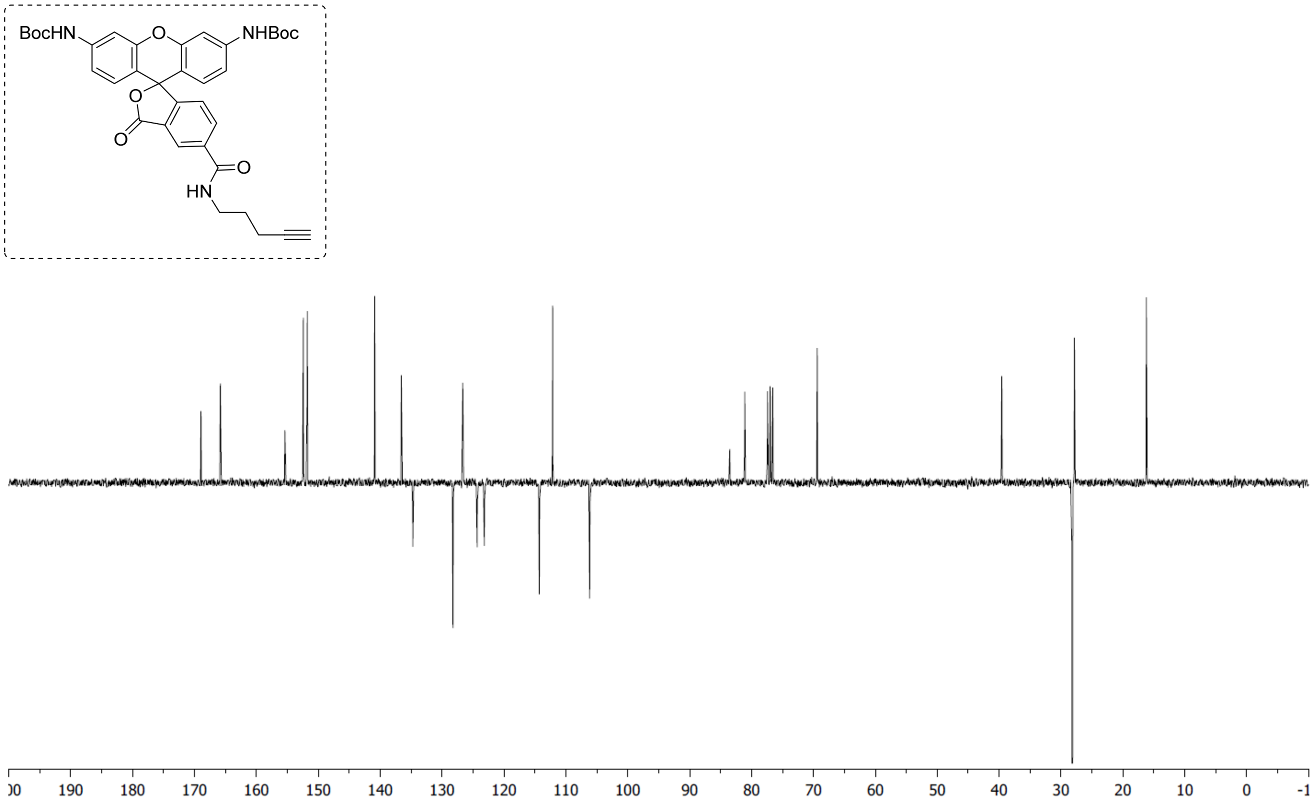


**Compound 6: $^1H$ NMR (300.17 MHz, MeOD)**

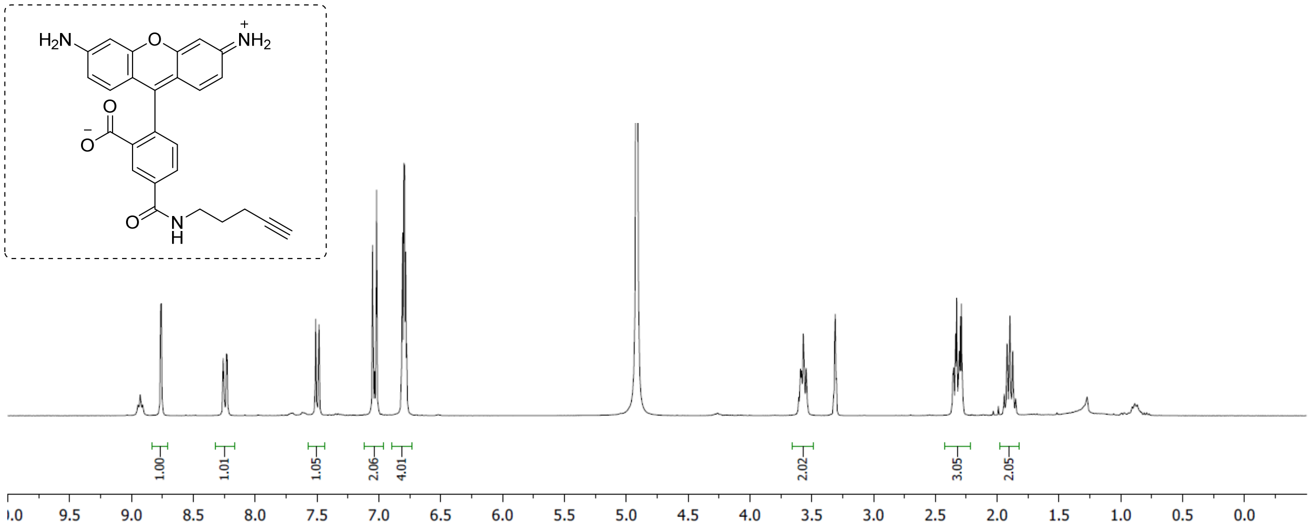

**Compound 6: $^{13}$C NMR (75.47 MHz, MeOD)**

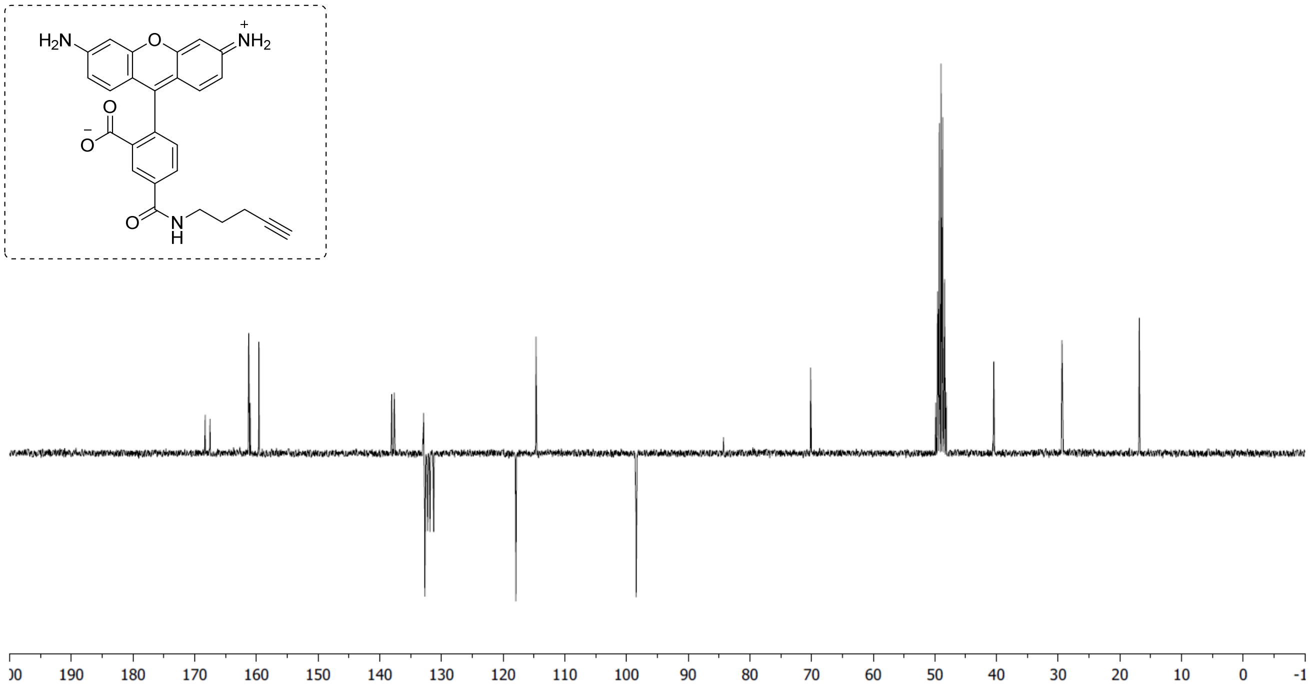

**Compound 9RK87: $^{1}$H NMR (300.17 MHz, DMSO)**

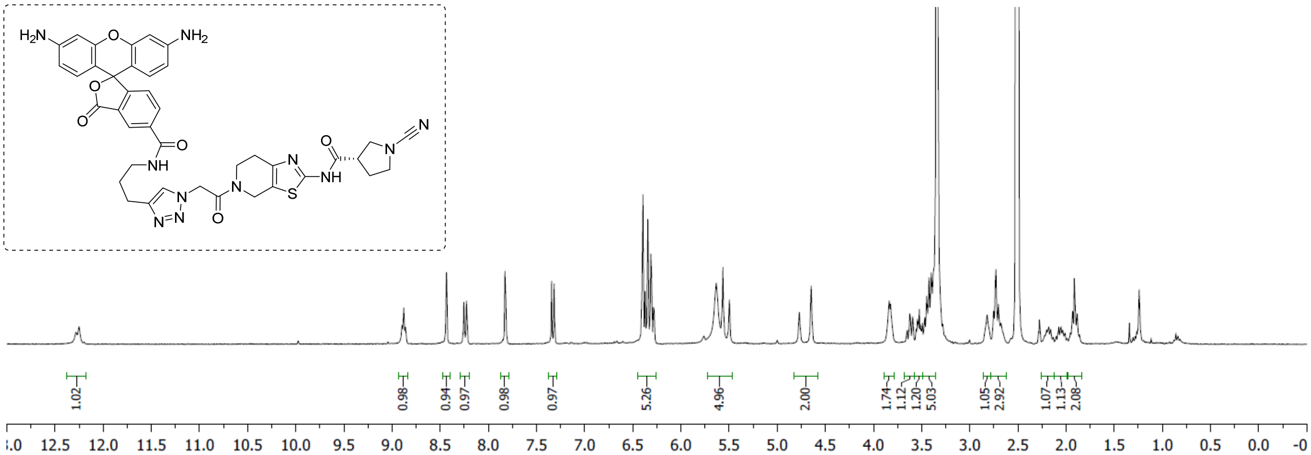

**Compound 7: $^{1}$H NMR (300.17 MHz, $CDCl_3$)**

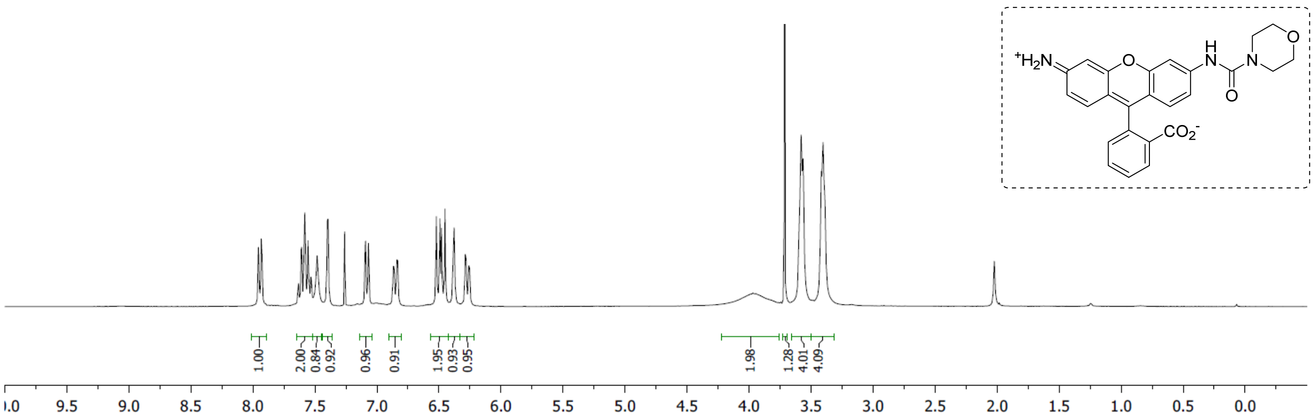

**Compound 7: $^{13}$C NMR (75.47 MHz, $CDCl_3$)**

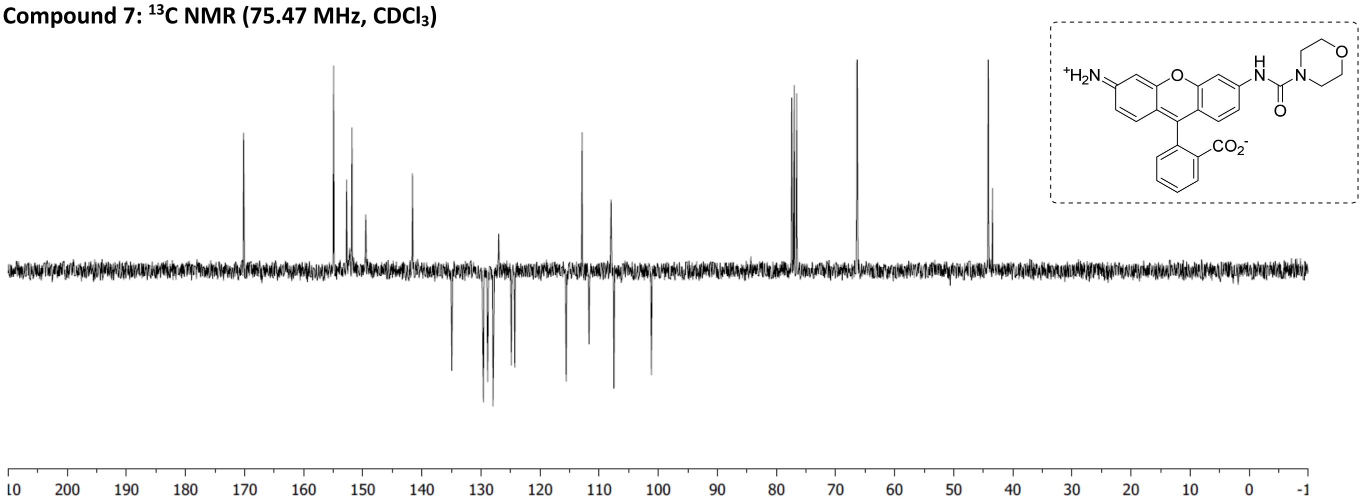


**Compound 8: $^{1}$H NMR (300.17 MHz, MeOD)**

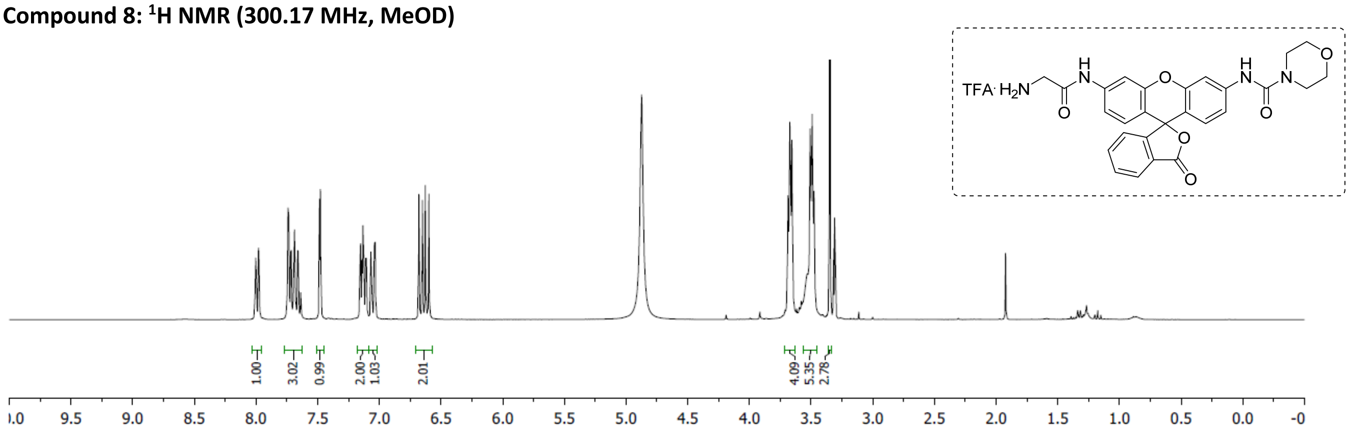


**Compound 8: $^{13}$C NMR (75.47 MHz, MeOD)**

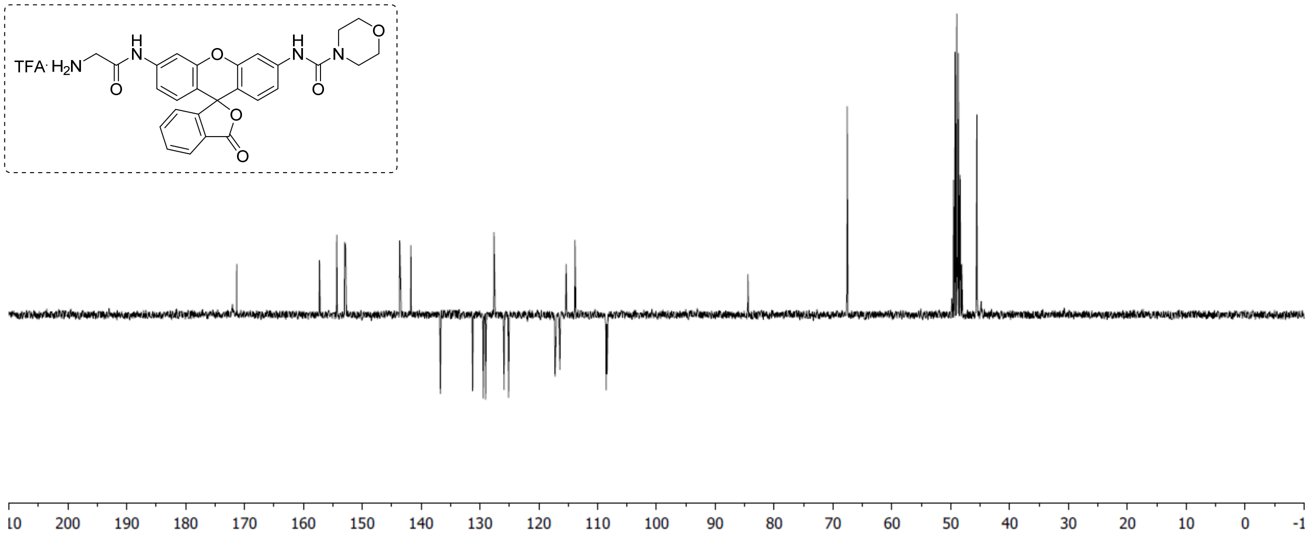

**Compound 8RK59: UV Trace.**

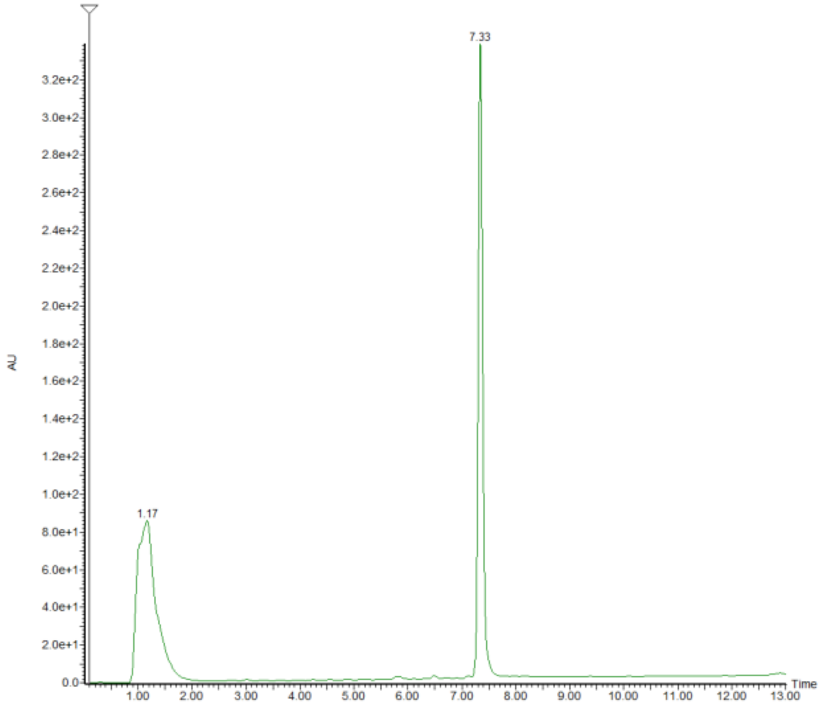


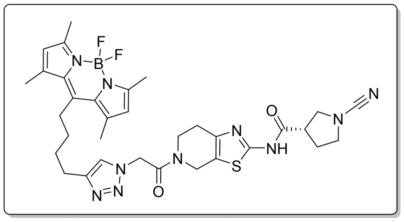

**Compound 9RK15: UV Trace.**

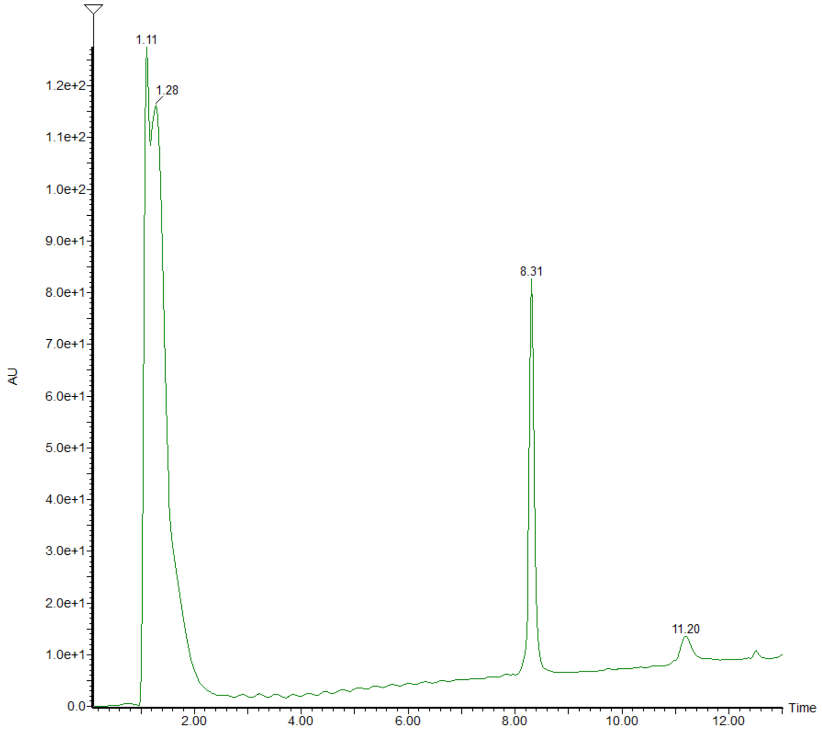


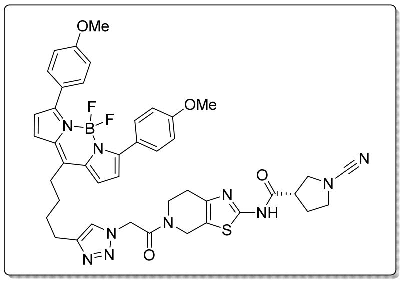

**Compound 9RK87: UV Trace.**

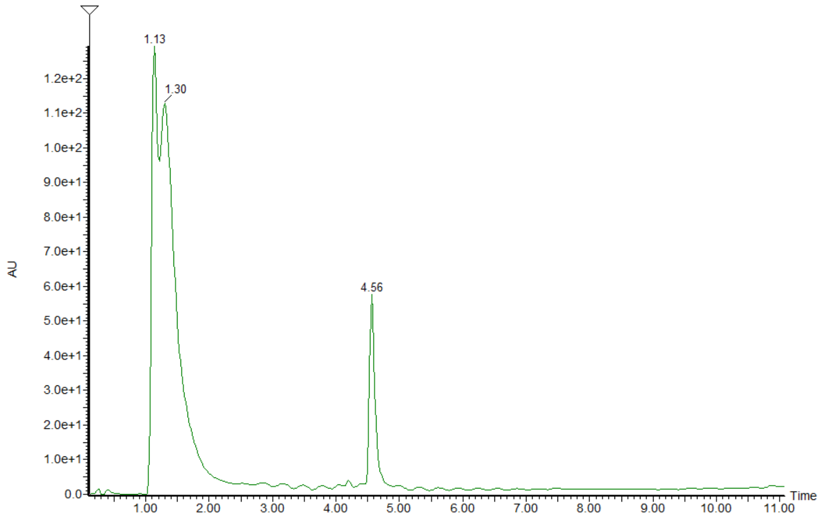


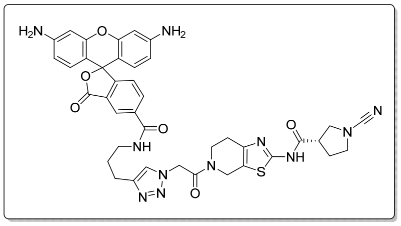


**Compound 11RK72: UV Trace.**

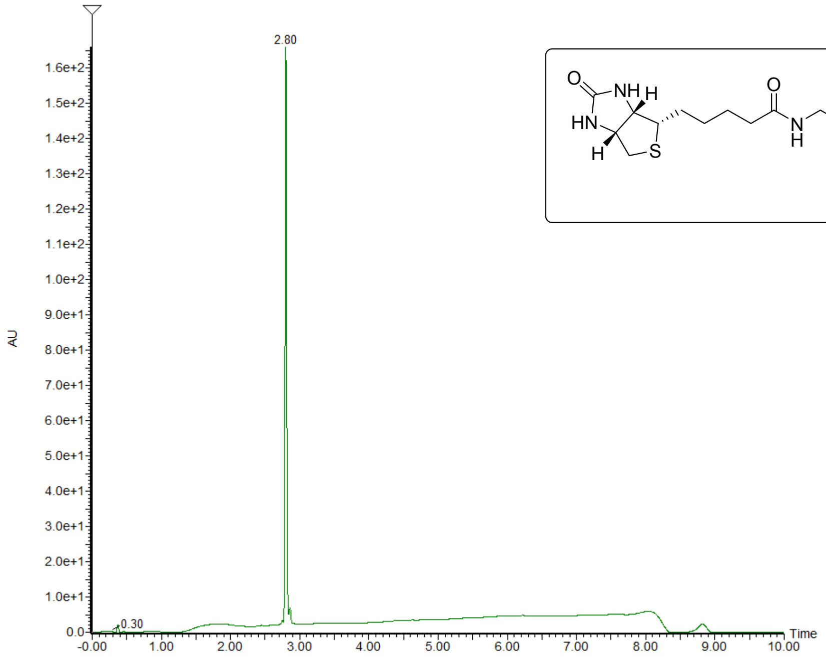


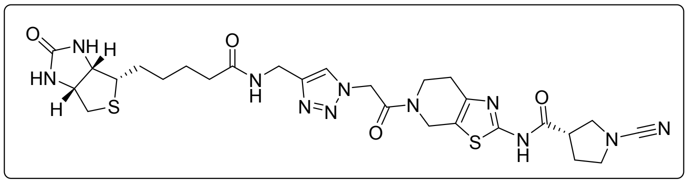

**Compound 11RK73: UV Trace.**

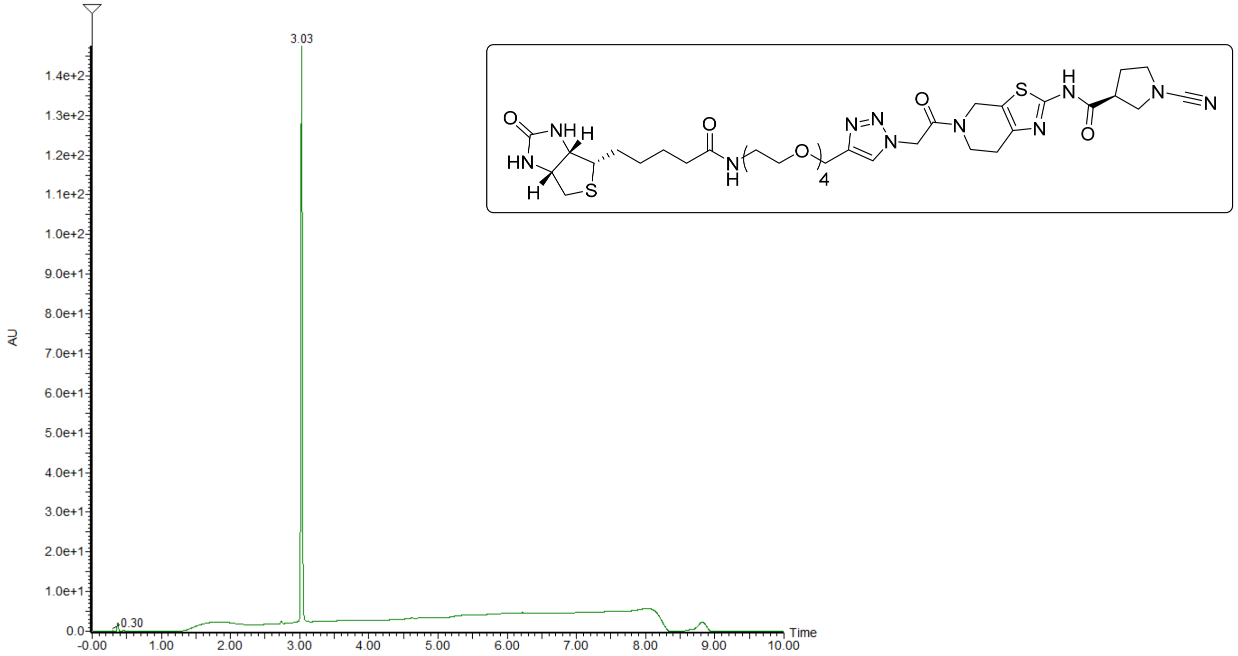


**Ub-Rho-morpholine LC-MS**

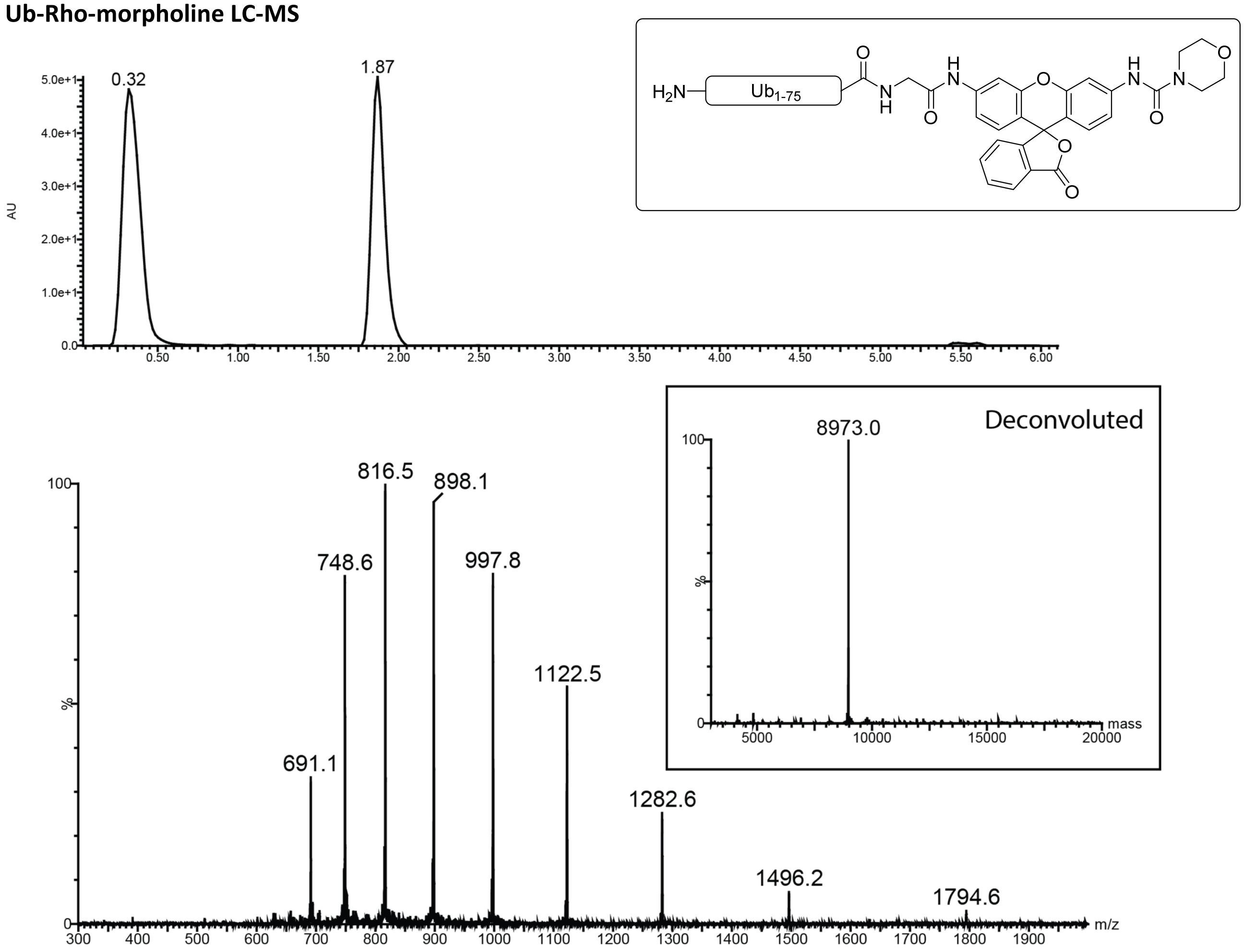
